# Transforming Growth Factor β1 Modulates Sex Differences in Cardiac Myofibroblast Activation on Hydrogel Biomaterials

**DOI:** 10.64898/2026.01.16.699818

**Authors:** Mason N. Faust, Ashley K. Nguyen, Rayyan M. Gorashi, Nicole E. Félix Vélez, Meaghan C. Loud, Brian A. Aguado

## Abstract

Cardiac fibrosis is a pathological process in which the myocardium stiffens due to the overproduction of extracellular matrix (ECM) proteins. Cardiac fibroblasts activate to myofibroblasts in response to the inflammatory cytokine transforming growth factor beta1 (TGF-β1) to promote fibrotic scarring. Biological sex also influences cardiac fibrosis progression and patient outcomes, where males exhibit increased fibrotic scarring after acute inflammation relative to females. At the cellular level, sex differences in TGF-β1-mediated cardiac myofibroblast activation processes have not been clearly defined. We hypothesized that TGF-β1 would cause sex-specific cardiac myofibroblast activation levels and alter the secretion of bioactive molecules to modulate sex differences in cardiac fibrosis. Primary left ventricle cardiac fibroblasts were isolated from male and female C57BL/6J mice and cultured on hydrogel biomaterials mimicking native myocardial ECM stiffness and treated with TGF-β1 and/or the TGF-β1 receptor inhibitor SD208. Male myofibroblasts exhibited increased α-SMA stress fiber formation, increased SMAD2/3 localization, and greater resistance to SD208 inhibition compared to female myofibroblasts on hydrogels at various time points tested. Sex differences in relative secreted cytokine abundance were also determined, with male CFs secreting increased vascular endothelial growth factor (VEGF) and female CFs producing increased periostin and fibroblast growth factor 21 in response to TGF-β1. Our findings establish that TGF-β1 mediates sex differences in cardiac myofibroblast activation on hydrogels and secreted factors that may modulate the myocardial microenvironment. Our work underscores the importance of using hydrogels as cell culture platforms to recapitulate sex-specific cardiac fibrosis phenotypes as a steppingstone towards identifying sex-dependent therapeutic interventions for cardiac fibrosis.

## Introduction

Cardiac fibrosis is a hallmark of multiple forms of cardiovascular disease, in which the myocardium stiffens due to the overproduction and buildup of extracellular matrix (ECM) proteins. Myocardial ECM stiffening disrupts the mechanical and electrochemical functions of a healthy heart, resulting in eventual heart failure for millions of patients worldwide^1–4^. Currently, there are no effective treatments to halt or reverse cardiac fibrosis once diagnosed, aside from candidate small molecule therapies that are used to slow progression and partially alleviate symptoms^5,6^. Furthermore, biological sex contributes substantially to biases in cardiac health and disease^7–10^. In one example of disease, female patients are three times more likely to develop heart failure with preserved ejection fraction (HFpEF) with increased concentric thickening and fibrotic stiffening of the left ventricle (LV) relative to males^11,12^. In another example, male patients are 3-4 times more likely to experience a myocardial infarct and have 1.25-fold higher rates of infarct expansion, scar tissue development, and neutrophil recruitment^13^. Furthermore, sex differences in baseline cardiac function have also been reported in rodents, which mirror observations in humans^14,15^. The context-specific sex differences apparent across multiple cardiovascular diseases and species have prompted us to improve our understanding of how biological sex contributes to cardiac fibrosis progression.

Cardiac fibroblasts (CFs) residing in the interstitial space of the myocardium are likely contributors to sex differences in myocardial fibrosis^16^. CFs activate to pathogenic myofibroblasts, which serve as the main producers of ECM proteins that subsequently accumulate and cause cardiac fibrosis^17–19^. Recent studies have demonstrated that male cardiac myofibroblasts have 2-fold increased production of fibrotic markers relative to female myofibroblasts when treated with either estradiol or isoproterenol to stimulate β-adrenergic receptor signaling, highlighting the need to understand sex-specific mechanisms of myofibroblast activation during cardiac fibrosis^20–22^. Cardiac myofibroblasts also secrete a myriad of inflammatory cytokines and matricellular proteins to recruit inflammatory immune cells in response to damaged myocardium^23,24^. In one example, cardiac myofibroblasts release the matricellular periostin to promote the adhesion and recruitment of inflammatory monocytes during the initiation of fibrosis^25^. In parallel, cardiac myofibroblasts also respond to secreted factors from inflammatory immune cells, causing a persistently activated myofibroblast phenotype that does not revert to a quiescent fibroblast^26^. Our collective understanding of sex differences in the cardiac myofibroblast activation and subsequent secretome during myocardial fibrosis progression must be improved to identify critical signaling pathways that can serve as future targets for sex-specific therapeutic interventions.

Transforming growth factor beta (TGF-β1) is likely the most well-documented secreted signaling molecule responsible for myofibroblast activation across multiple tissues^27^. TGF-β1 exerts its effects through the activation of a family of transcription factors in fibroblasts called small mothers against decapentaplegic (SMAD) proteins, which have been shown to directly promote alpha smooth muscle actin (α-SMA) expression^28,29^. Previous work on CFs suggests that TGF-β1 and SMAD-mediated gene expression also have a direct influence on the CF secretome^30–32^. Despite these advances, it remains unknown whether TGF-β1 regulates the CF secretome in a sex-dependent manner. Previous studies have also shown that circulating androgens in male mice contribute to increased expression of TGF-β1 during myocardial remodeling in response to pressure overload^33^, suggesting sex-specific mechanisms in TGF-β1 signaling regulation. Furthermore, small molecule drugs such as SD208 have been used as TGFBR1 inhibitors to deactivate myofibroblasts via reduced TGF-β1 signaling^34,35^, but their sex-specific impact on cardiac myofibroblast phenotypes have not yet been explored. As such, improving our understanding on how TGF-β1 causes sex differences in cardiac myofibroblast activation and how biological sex contributes to TGFBR1 inhibitor responses is warranted.

Investigating sex differences in mechanically sensitive CFs also require appropriately bioengineered cell culture platforms to recapitulate the myocardial ECM. CFs removed from the native myocardial ECM and plated on traditional tissue culture polystyrene (TCPS) cause the rampant activation of cardiac myofibroblasts *in vitro* via increased α-SMA expression^36–38^. Hydrogels have been widely used as cell culture platforms because of their ability to be specifically tuned to mimic the matrix stiffness found in the resident tissue^39^. For example, previous work has observed sex differences in valvular interstitial cells^40–44^ and endothelial cells^45,46^ cultured on hydrogels, relative to TCPS cultures that reveal minimal sex differences in phenotypes. Previous work has also established synergistic effects between hydrogel stiffness and TGF-β1 dosing with CF phenotypes^31^, but sex-dependent effects on the secretome have yet to be explored. Hydrogels may also serve as valuable precision biomaterial tools^47,48^ to determine sex differences in cardiac myofibroblast phenotypes and decouple the effects of mechanical cues from biochemical factors such as TGF-β1.

Here, we harness hydrogel cell culture platforms to investigate the role of TGF-β1 in regulating sex differences in cardiac myofibroblast activation and subsequent secretome profiles. First, we cultured male and female mouse CFs with TGF-β1 and evaluated the sex-specific response in myofibroblast activation and secretome. We also tested the effects of the antifibrotic drug SD208 to identify sex-specific responses to the inhibition of TGF-β1 signaling via TGFBR1 in cardiac myofibroblasts. After understanding the intracellular effects of TGF-β1 signaling, we characterized sex-dependent differences in the cardiac myofibroblast secretome. Collectively, our work identifies key sex differences in how CFs respond to TGF-β1 as a cell culture input and provides insights into the biological processes that give rise to sex-dependent myocardial fibrosis.

## Results

### Hydrogels maintain male and female cardiac fibroblast quiescence in vitro

We first sought to engineer a hydrogel culture platform that mimics the mechanical properties of the healthy myocardium to preserve the quiescent CF phenotype *in vitro*. The supraphysiologic stiffness of tissue culture polystyrene (TCPS) has an elastic modulus (E ∼ 1 GPa) that far exceeds the stiffness of the native mouse myocardium (E ∼ 20 kPa)^49,50^. Recognizing that CFs are mechanically sensitive cells and aberrantly activate to myofibroblasts when cultured on TCPS^38^, we implemented a poly(ethylene glycol) (PEG) hydrogel cell culture platform that recapitulates the native stiffness of healthy myocardium (**Figure 1A**). Leveraging thiol-ene click chemistry^51^, we synthesized a norbornene-functionalized 8-arm PEG polymer designed to crosslink with a PEG dithiol linker and a cysteine-bearing CRGDS adhesive peptide (**Figure 1B**). We initiated hydrogel polymerization under UV light (365 nm) and using oscillatory shear rheology to measure the storage and loss moduli during crosslinking, we calculated the Young’s modulus of the hydrogels to be 24.1 ± 0.6 kPa (**Figure 1C**).

**Fig. 1:**
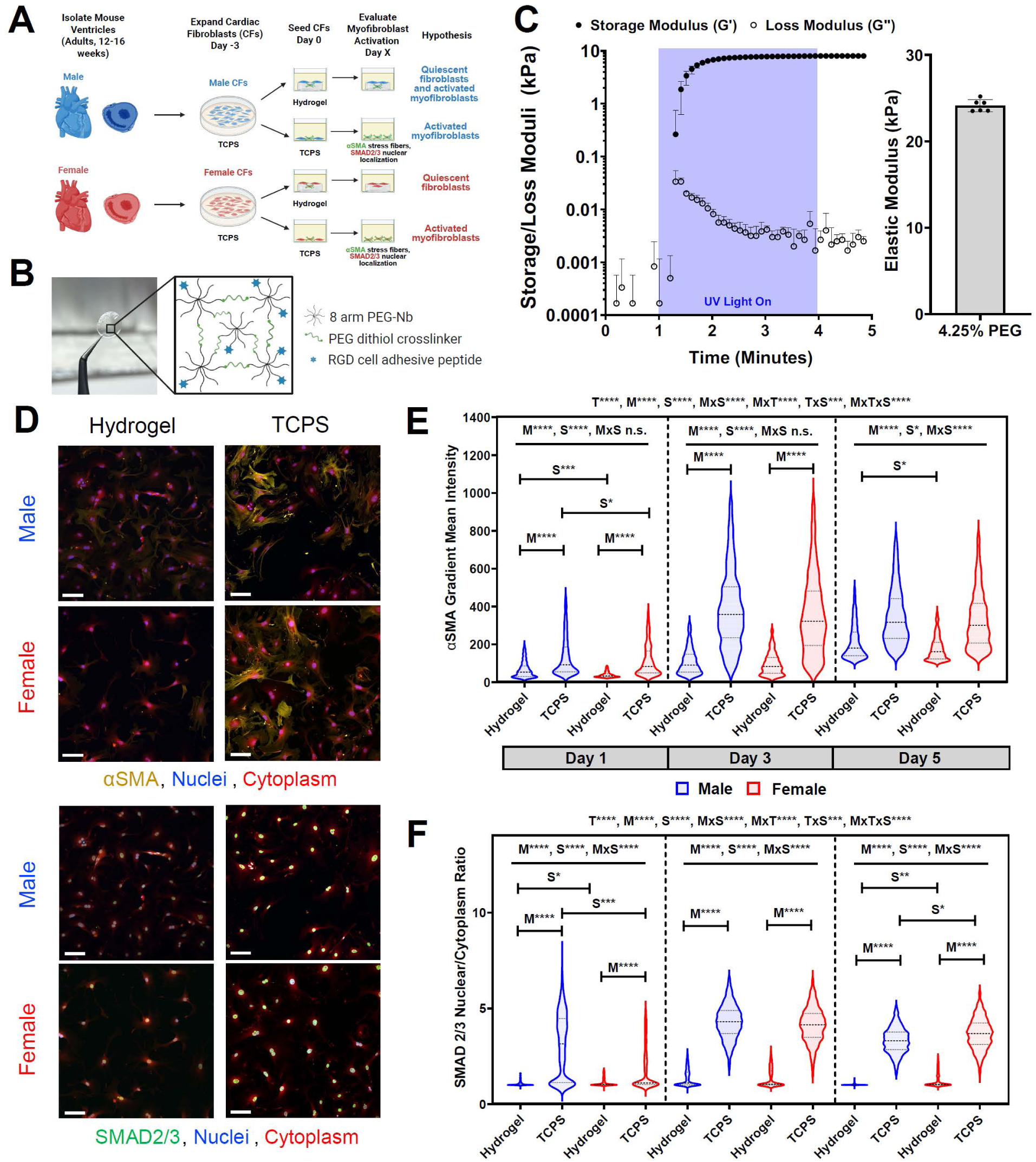
PEG hydrogels maintain male and female cardiac fibroblast quiescence relative to TCPS. (**A**) Schematic describing experimental design including proposed hypotheses for male and female CF phenotypes on PEG hydrogel and TCPS culture platforms. (**B**) Schematic of hydrogel components and photograph of a hydrogel on a 12 mm coverslip for cell culture. (**C**) Rheological characterization of UV-mediated polymerization of PEG-Nb crosslinked with PEG-dithiol. Elastic modulus of hydrogel calculated from the final plateau storage modulus measurement (4.25% PEG formulation). N = 6 hydrogels, mean ± standard deviation (S.D.) shown. (**D**) Representative images of male and female CFs cultured on hydrogels and TCPS (Day 3 time point shown). Scale bars = 100 μm. (**E-F**) Quantification of (**E**) α-SMA gradient mean intensity and (**F**) nuclear localization of SMAD2/3 in male and female CFs cultured on hydrogels and TCPS at various time points. N=3 hydrogels, mean ± S.D. shown. Significance was determined using (1) three-way nonparametric ANOVA (S=sex, M=material, T=time, x=interaction effects between variables) and (2) two-way nonparametric ANOVA (P < 0.001) and a Cohen’s d test (*d < 0.2, **d < 0.5, ***d < 0.8, and ****d < 1.4) denoting significance between sex.

We next cultured primary CFs isolated from healthy mouse hearts on our hydrogels, with the hypothesis that hydrogel culture would circumvent the rampant myofibroblast activation observed on TCPS. We also hypothesized mouse male CFs would exhibit elevated myofibroblast activation relative to female CFs on hydrogels, given prior observations in rat CFs^20^. We assessed the effects of male and female CF culture on hydrogel vs. TCPS over 5 days by evaluating α-SMA stress fiber formation and SMAD2/3 nuclear localization as outputs for myofibroblast activation (**Figure 1D**). We first quantified the gradient mean intensity of immunostained α-SMA for stress fiber formation and observed a significant reduction in myofibroblast activation on hydrogel relative to TCPS across all time points tested (**Figure 1E**). Statistical analysis showed that Time (T), Sex (S), and Material (M), along with all possible interactions between variables, significantly impacted α-SMA stress fiber formation (P < 0.0001). Post-hoc tests showed that at Day 1, we observed a ∼2-fold reduction in male cardiac myofibroblast activation and a ∼3-fold reduction in female cardiac myofibroblast activation on hydrogels relative to TCPS. As expected with increased culture time on TCPS, we observed a significant increase in myofibroblast activation on TCPS at Days 3 and 5 relative to Day 1. On hydrogels, α-SMA expression remained relatively constant across all time points, indicating minimal myofibroblast activation. We confirmed hydrogel culture was able to inhibit myofibroblast activation for both males and females across all time points, with a ∼3.5-fold and ∼3-fold reduction in male α-SMA expression and a ∼3.75-fold and ∼2.5-fold reduction in female α-SMA on hydrogels relative to TCPS at days 3 and 5, respectively. Additionally, male CFs had ∼1.7-fold and ∼1.2-fold significantly higher activation on hydrogels at Day 1 and Day 5 relative to female CFs, supporting our initial hypothesis.

We also characterized SMAD2/3 nuclear localization in male and female CFs on hydrogel and TCPS culture substrates, where increased nuclear localization is linked to myofibroblast activation. We observed reduced SMAD2/3 nuclear localization on hydrogels relative to TCPS across all time points tested, trending similarly to the myofibroblast activation previously observed (**Figure 1D,F**). Nuclear localization was quantified as the ratio of the nucleus to cytoplasmic expression of immunostained SMAD2/3. Similar to α-SMA, SMAD2/3 nuclear localization was significantly impacted by Time (T), Sex (S), and Material (M), along with all possible interactions between variables. At Day 1, we observed a ∼3-fold lower nuclear localization in male SMAD2/3 expression and a ∼1.5-fold lower nuclear localization in female SMAD2/3 expression on hydrogels relative to TCPS. At Days 3 and 5 in culture, we also observed a significant increase in SMAD2/3 nuclear localization on TCPS, whereas lower SMAD2/3 nuclear localization on hydrogels remained relatively constant across all time points, further suggesting culture on hydrogels inhibits myofibroblast activation. CF area was also quantified on hydrogels and TCPS, acknowledging that myofibroblasts have larger cell areas^38^. CFs in culture have increased cell area on TCPS relative to hydrogels, with a significant increase in cell area observed in female CFs relative to male CFs at Day 3 (**Supplementary Figure 1**).

### Sex differences in TGF-β-induced cardiac myofibroblast activation on hydrogels

To more robustly probe potential differences in myofibroblast activation between male and female CFs, we induced myofibroblast activation in CFs via TGF-β1 stimulation (**Figure 2A**). Again, we assessed myofibroblast activation using α-SMA stress fiber formation and SMAD2/3 nuclear localization. We first supplemented TGF-β1 into the culture media at physiologically relevant concentrations^52,53^ of 1 and 10 ng/mL to determine the concentration that would yield the greatest increase in myofibroblast activation on hydrogels. As expected, we observed increased α-SMA expression in both male and female CFs after 3 days of TGF-β1 treatment (**Figure 2B**). At a dose of 1 ng/mL, female CFs had a greater increase in α-SMA expression relative to male CFs. However, at a higher TGF-β1 dose of 10 ng/mL, we observed male CFs to have a ∼4-fold increase in α-SMA expression relative to female CFs (**Figure 2C**). Interestingly, we observed similar α-SMA expression in female CFs at both 1 and 10 ng/mL TGF-β1, suggesting less sensitivity to TGF-β1 dosing concentration in female cells. Female CFs had significantly higher cell area relative to male CF after 1 ng/mL TGF-β1 treatment but had equal cell areas with and without 10 ng/mL TGF-β1 treatment (**Supplementary Figure 2A**). Taken together, these data suggest that while female CFs show increased activation at lower TGF-β1 concentrations, male CFs are more responsive to TGF-β1 treatment relative to female CFs, particularly at higher doses of TGF-β1.

**Fig. 2:**
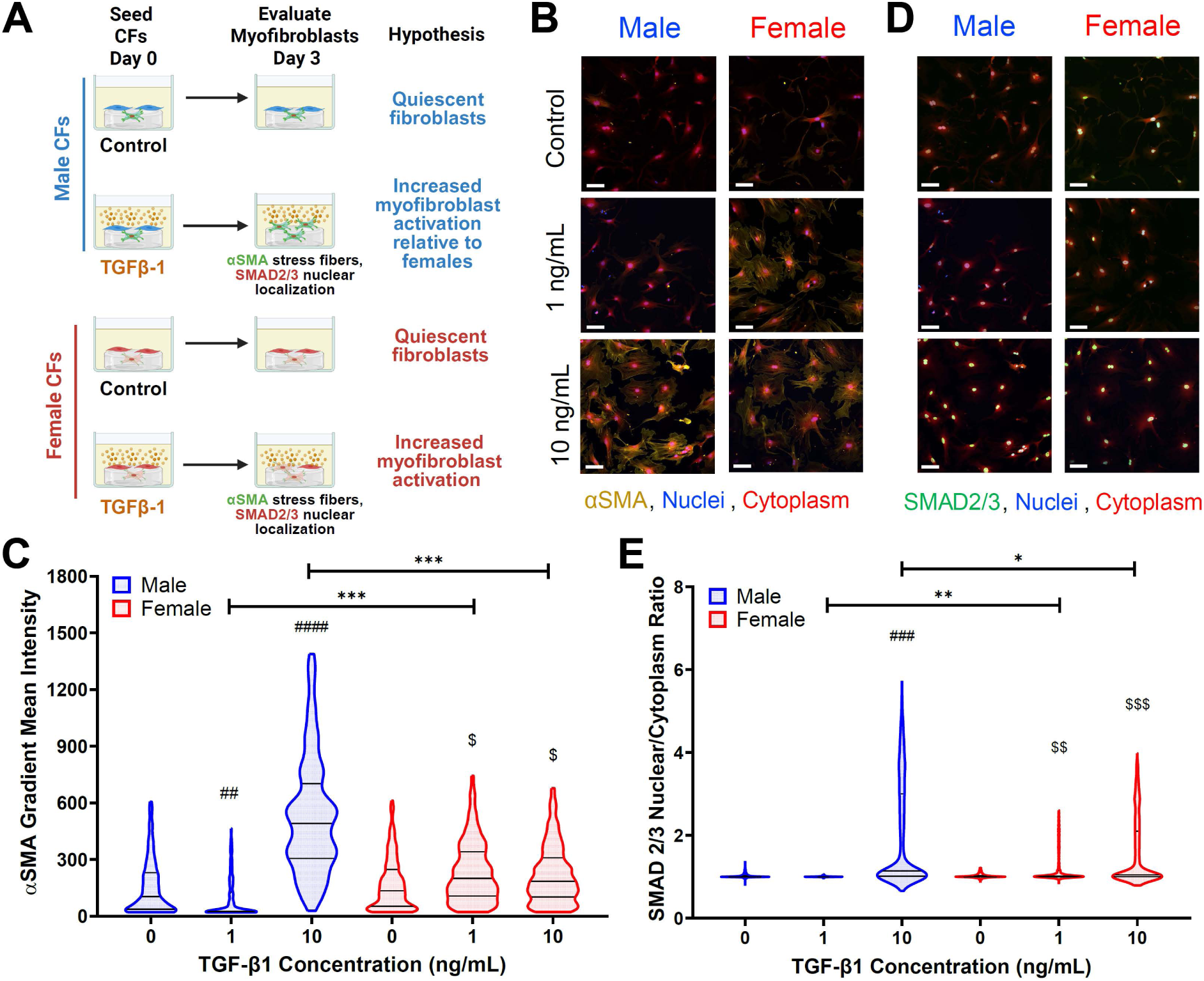
TGF-β1 drives sex differences in cardiac myofibroblast activation and SMAD2/3 nuclear localization on hydrogels. (**A**) Schematic describing experimental design including proposed hypotheses for sex differences in cardiac myofibroblast phenotypes treated with TGF-β1 on hydrogels. (**B-C**) Representative images and quantified α-SMA gradient mean intensity values for male and female CFs cultured on hydrogels and treated with 0, 1, and 10 ng/mL TGF-β1. Scale bar = 100 μm. N=3 hydrogels, n>432 cells, mean ± S.D. shown. (**D-E**) Representative images and quantified SMAD2/3 nuclear to cytoplasmic intensity ratios for male and female CFs cultured on hydrogels and treated with 0, 1, and 10 ng/mL TGF-β1. Scale bar = 100 μm. N=3 hydrogels, n>806 cells, mean ± S.D. shown. Significance for (**C)** and (**E**) was determined using a two-way nonparametric ANOVA data (P < 0.001) with a Cohen’s d test (*d < 0.2, **d < 0.5, ***d < 0.8, and ****d < 1.4) denoting significance between sex; hashtag indicates significance in male group (^#^d < 0.2, ^##^d < 0.5, ^###^d < 0.8, and ^####^d < 1.4) denoting significance relative to male control; dollar sign indicates significance in female group (^$^d < 0.2, ^$$^d < 0.5, ^$$$^d < 0.8, and ^$$$$^d < 1.4) denoting significance relative to female control.

We also hypothesized male CFs would have increased nuclear localization of SMAD2/3 relative to female CFs, as SMAD2/3 are transcription factors that translocate from the cytoplasm to the nucleus in the presence of TGF-β1^29^ (**Figure 2A**). Consistent with our α-SMA stress fiber formation results, we observed increased female nuclear localization of SMAD2/3 relative to male CFs when treated with 1 ng/mL of TGF-β1 (**Figure 2D-E**). When the dose was increased to 10 ng/mL, we observed a greater increase of SMAD2/3 nuclear localization in male CFs relative to female CFs, again suggesting increased myofibroblast activation in males relative to females at higher doses of TGF-β1. Together, we observe sex differences in myofibroblast activation based on α-SMA expression and SMAD2/3 nuclear localization as a function of TGF-β1 treatment on hydrogels.

### Sex differences in cardiac myofibroblast responses to SD208 inhibition on hydrogels

Given the heterogeneity in myofibroblast activation levels observed in male and female CFs on hydrogels, we next sought to determine any sex differences in cardiac myofibroblast responses to SD208, a TGFBR1 kinase receptor inhibitor. As an inhibitor of TGF-β1, we hypothesized that SD208 would reduce α-SMA stress fiber formation and SMAD2/3 nuclear localization in CFs, and female CFs would be more responsive to the inhibitor based on other studies in aortic valve myofibroblasts^43^ (**Figure 3A**). We cultured male and female CFs dosed with SD208 in the culture medium at concentrations of 1, 10, and 100 μM for 3 days to deactivate myofibroblasts. As expected, α-SMA expression was significantly reduced in both males and female CFs (**Figure 3B**). In male CF cultures, we observed decreases in α-SMA expression that correlated with increasing SD208 concentrations. In contrast, we observed that female CF α-SMA expression decreased significantly at just 1 μM SD208 and plateaued even when higher concentrations of SD208 were used (**Figure 3C**). Interestingly, we observed male CFs to have increased SMAD2/3 nuclear localization as a function of increasing concentration of SD208, whereas female CFs had no change in SMAD2/3 nuclear localization with increasing concentration (**Figures 3D-E**). Cell areas were consistent across SD208 treatment groups, indicating no major reductions in cell viability or increased apoptosis (**Supplementary Figure 2B**). In summary, SD208 elicits sex-dependent effects on α-SMA expression and SMAD2/3 nuclear localization when CFs are cultured on hydrogels.

**Fig. 3:**
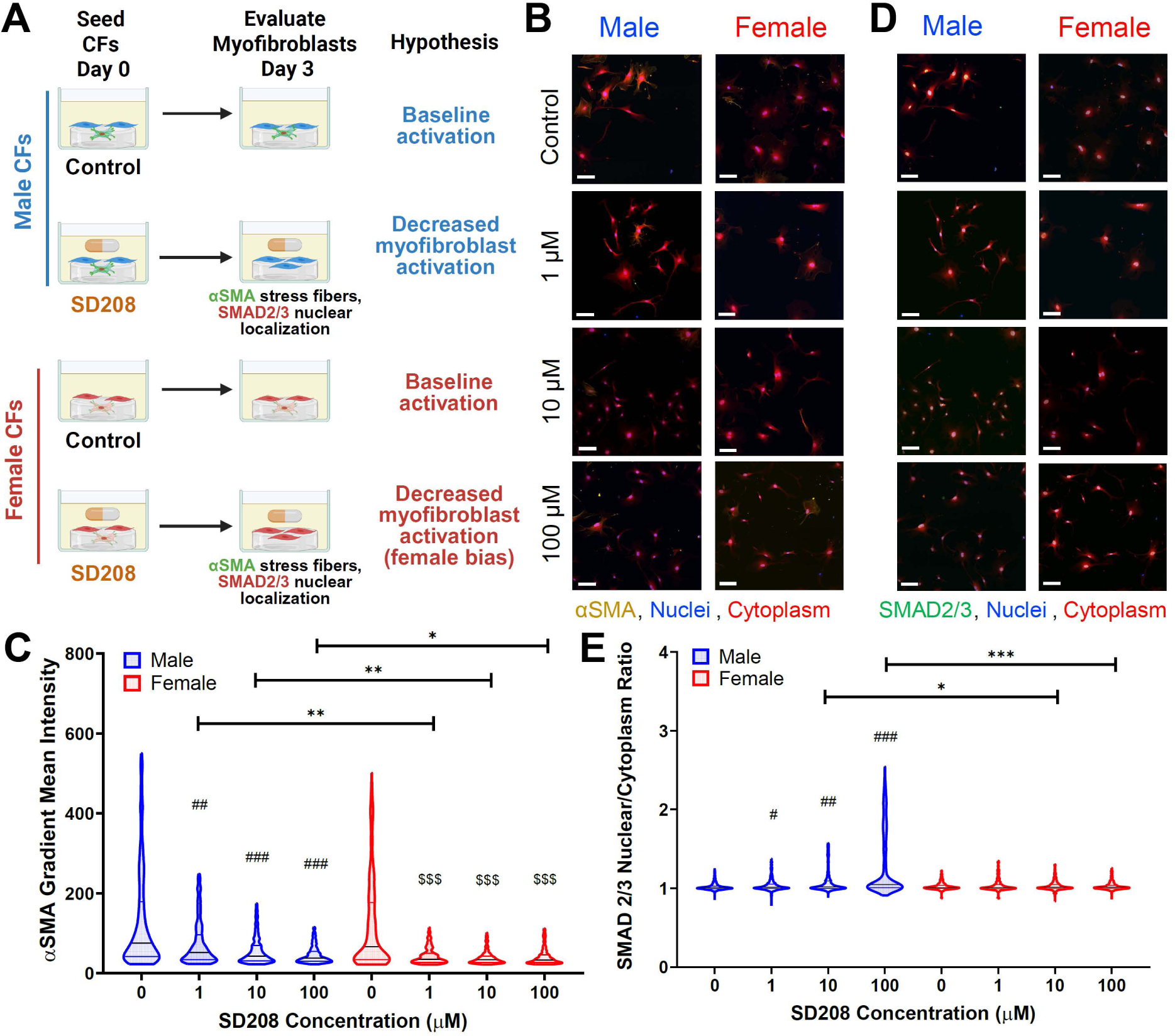
SD208 modulates sex differences in cardiac myofibroblast deactivation on hydrogels. (**A**) Schematic describing experimental design including proposed hypotheses for sex differences in cardiac myofibroblast phenotypes treated with SD208 on hydrogels. (**B-C**) Representative images and quantified α-SMA gradient mean intensity values for male and female CFs cultured on hydrogels and treated with 0, 1, 10, and 100 μM of SD208. Scale bar = 100 μm. N=3 hydrogels, n>90 cells, mean ± S.D. shown. (**D-E**) Representative images and quantified SMAD2/3 nuclear to cytoplasmic intensity ratios for male and female CFs cultured on hydrogels and treated with treated with 0, 1, 10, and 100 μM of SD208. Scale bar = 100 μm. N=3 hydrogels, n>263 cells, mean ± S.D. shown. Significance for (**C**) and (**E**) was determined using a two-way nonparametric ANOVA data (P < 0.001) with a Cohen’s d test (*d < 0.2, **d < 0.5, ***d < 0.8, and ****d < 1.4) denoting significance between sex; hashtag indicates significance in male group (^#^d < 0.2, ^##^d < 0.5, ^###^d < 0.8, and ^####^d < 1.4) denoting significance relative to male control; dollar sign indicates significance in female group (^$^d < 0.2, ^$$^d < 0.5, ^$$$^d < 0.8, and ^$$$$^d < 1.4) denoting significance relative to female control.

### SD208 modulates sex differences in cardiac myofibroblast phenotypes in the presence of TGF-β1

Next, we evaluated whether SD208 could suppress myofibroblast activation in male and female CFs in the continued presence of TGF-β1. CFs on hydrogels were treated with optimized doses of TGF-β1 (10 ng/mL), SD208 (1 μM), or both for 3 days. Since we observed male CFs are more responsive to TGF-β1, while female CFs are more sensitive to SD208, we hypothesized that combined treatment would differentially modulate α-SMA stress fiber formation and SMAD2/3 nuclear localization between the sexes. Consistent with earlier findings, we confirmed that TGF-β1 increased α-SMA expression in male CFs relative to female CFs, whereas SD208 reduced α-SMA expression more robustly in female CFs (**Figures 4A-B**). When SD208-treated cultures were simultaneously treated with TGF-β1, myofibroblast activation was partially restored in both sexes, but more robustly in female CFs (∼2.1-fold increase over SD208 alone) than in males (∼1.3-fold increase), indicating that female CFs retain a greater capacity to maintain TGF-β1 driven activation regardless of exposure to SD208.

**Fig. 4:**
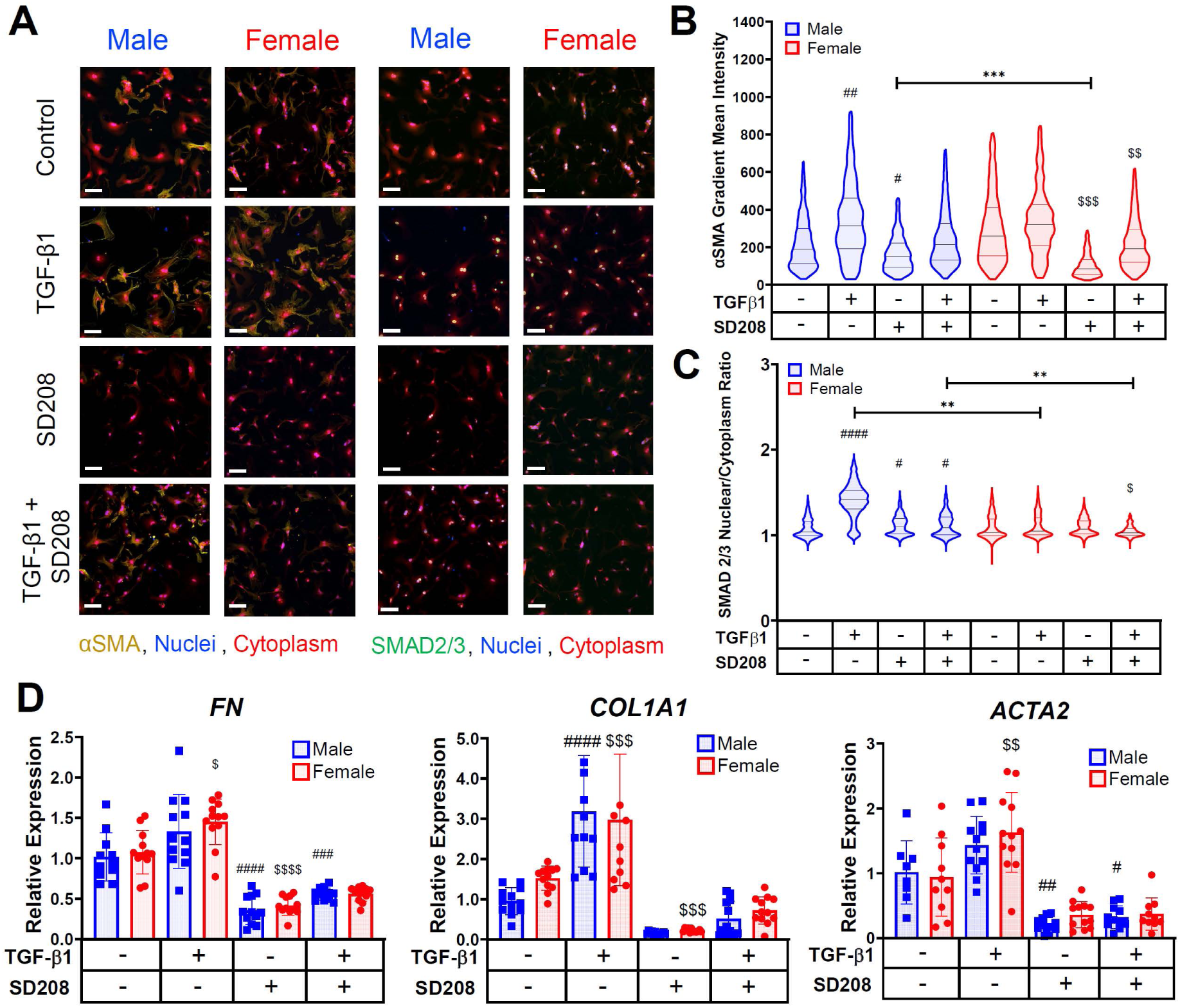
Sex differences in cardiac myofibroblast responses to combined SD208 and TGF-β1 treatments on hydrogels. (**A**) Representative images of male and female CFs cultured on hydrogels with TGF-β1 and/or SD208. Scale bar = 100 μm. (**B**) Quantified α-SMA gradient mean intensity values for male and female CFs cultured on hydrogels with TGF-β1 and/or SD208. (**C**) SMAD2/3 nuclear localization intensity ratios for male and female CFs cultured on hydrogels with TGF-β1 and/or SD208. Significance for (**B)** and (**C)** was determined using a two-way nonparametric ANOVA data (P < 0.001) with a Cohen’s d test (*d < 0.2, **d < 0.5, ***d < 0.8, and ****d < 1.4) denoting significance between sex; hashtag indicates significance in male group (^#^d < 0.2, ^##^d < 0.5, ^###^d < 0.8, and ^####^d < 1.4) denoting significance relative to male control; dollar sign indicates significance in female group (^$^d < 0.2, ^$$^d < 0.5, ^$$$^d < 0.8, and ^$$$$^d < 1.4) denoting significance relative to female control. (**D**) RT-qPCR gene expression measurement of *FN*, *COL1A1,* and *ACTA2* in CFs cultured on hydrogels treated with TGF-β1 and/or SD208. Values were normalized to male hydrogel control (N=12, mean ± S.D. shown, two-way ANOVA, *P≤0.05, **P≤0.01, ***P≤0.001, ****P≤0.0001).

We also quantified SMAD2/3 nuclear localization in the presence of both TGF-β1 and SD208 (**Figure 4C**). In males, TGF-β1 alone increased SMAD2/3 nuclear localization, which was reduced with SD208 treatment. Notably, the combination of TGF-β1 and SD208 did not reinstate SMAD2/3 nuclear localization in male CFs, suggesting sustained inhibition of upstream TGF-β1 signaling. In female CFs, neither TGF-β1, SD208, nor their combination increased SMAD2/3 nuclear localization, with dual treatment resulting in lower SMAD2/3 nuclear localization than in males, corroborating our prior results (**Figure 3B**). Cell areas were consistent across groups, except for increased female CF area in the TGF-β1 treatment group, and increased cell area in the male CF group treated with both TGF-β1 and SD208 (**Supplementary Figure 3**).

Finally, real time quantitative polymerase chain reaction (RT-qPCR) analysis showed that TGF-β1 increased expression of *FN*, *COL1A1*, and *ACTA2* in both male and female CFs on hydrogels, whereas SD208 alone reduced expression of all three myofibroblast genes (**Figure 4D**). Adding TGF-β1 in the presence of SD208 did not rescue gene expression of *FN, COL1A1*, and *ACTA2* in neither male nor female CFs, even though we initially observed rescue in α-SMA stress fiber formation at the protein level (**Figure 4B**). Our data suggest rescue of α-SMA stress fiber formation occurs downstream of transcription and that this regulation may be sex-biased. Together, these data demonstrate that SD208 mitigates TGF-β1-driven myofibroblast activation on hydrogels, with males exhibiting stronger SMAD2/3 activation and females showing greater α-SMA stress fiber development during reactivation. These sex-specific differences in sensitivity to TGF-β1 and SD208 are suggestive of differing fibrotic responses between male and female CFs on substrates with physiologically relevant stiffness.

### TGF-β1 modulates sex differences in the cardiac myofibroblast secretome on engineered hydrogels

After establishing that TGF-β1 signaling differentially regulates myofibroblast-associated gene and protein expression in male and female CFs, we next asked whether their secretomes were similarly sex-biased. We collected conditioned media from male and female CFs cultured on hydrogels with or without TGF-β1 and measured the abundance values for 111 mouse inflammatory cytokines (**Supplemental Figure 4, Supplementary Table 2**). Unsupervised hierarchical clustering of differentially expressed inflammatory cytokines in conditioned media, based on correlation distance and average linkage, indicated coordinated, sex-dependent changes in secreted factors based on biological sex and TGF-β1 treatment (**Figure 5A**). Within each sex, TGF-β1 treatment altered the abundance of 11 cytokines in males (8 upregulated and 3 downregulated), and 13 in females (9 upregulated and 4 downregulated) relative to untreated controls (**Figures 5B-C**), demonstrating TGF-β1-associated changes in secretory profiles for both males and females.

**Fig. 5:**
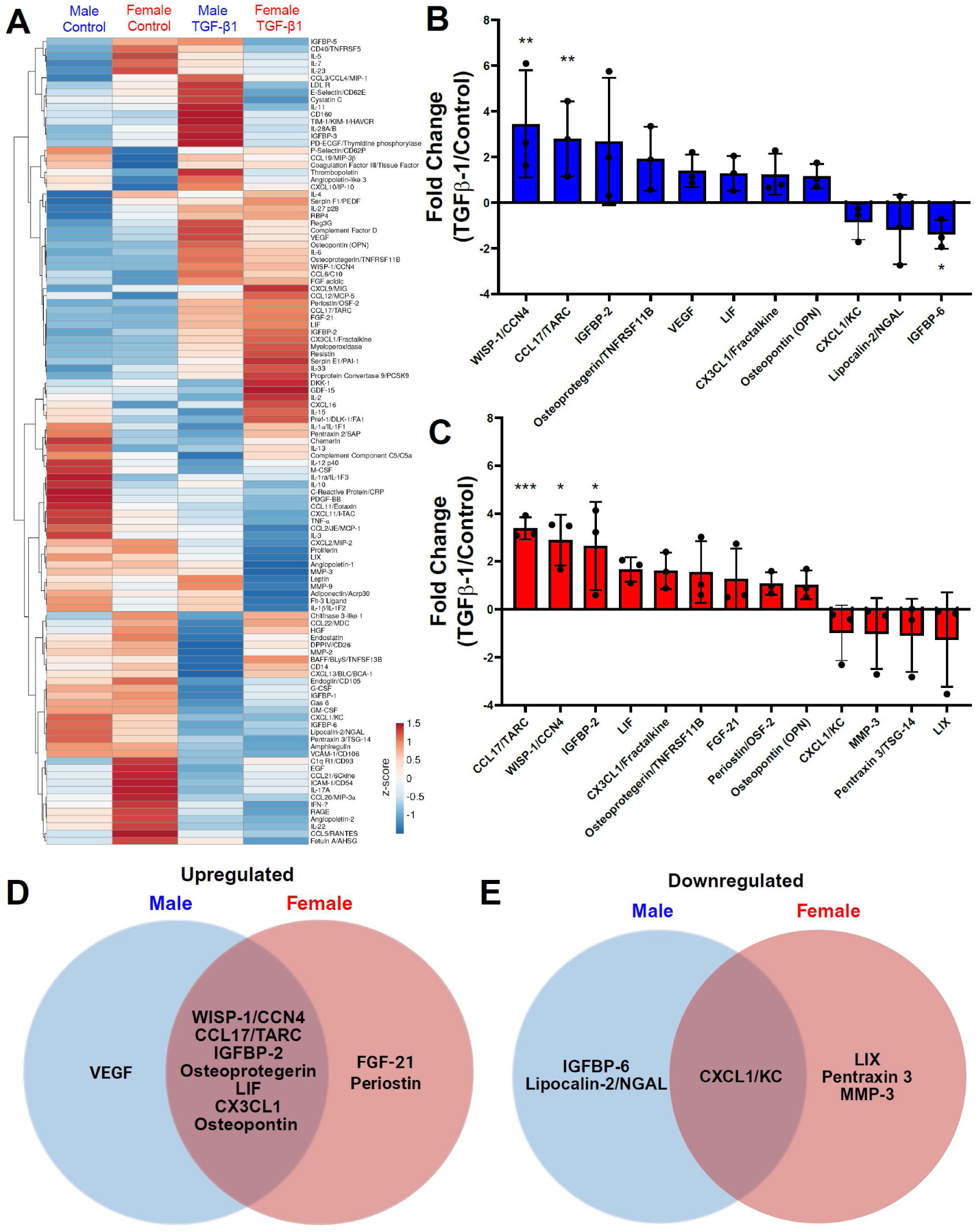
TGF-β1 treatment induces sex differences in the cardiac fibroblast secretome on engineered hydrogels. **(A)** Heat map showing log2 fold changes in secreted cytokine abundances in male and female CFs cultured with or without TGF-β1 using unit variance scaling. **(B-C)** Fold change graphs showing up- and down-regulated cytokines secreted from TGF-β1-treated CFs normalized to untreated CFs (Log2FC, cutoff ±1) for **(B)** male and (**C**) female CFs. **(D-E)** Venn diagrams comparing (**D**) upregulated and (**E**) downregulated cytokine secretion of male and female CFs treated with TGF-β1. Significance in fold change determined with Student t-tests for individual proteins (*P≤0.05, **P≤0.01, ***P≤0.001). N = 3 biological replicates, mean ± S.D. shown.

Male and female fibroblasts also shared a core TGF-β1 response, with Wnt1-Inducible Signalling Pathway protein 1 (WISP-1), C-C motif Chemokine Ligand 17 (CCL17), Insulin Growth Factor Binding Protein 2 (IGFBP2), Osteoprotegerin, Leukemia Inhibitory Factor (LIF), chemokine C-X3-C motif Ligand 1 (CX3CL1), and Osteopontin commonly upregulated, and C-X-C-motif Chemokine Ligand 1 (CXCL1) commonly downregulated (**Figures 5D-5E**). Interestingly, FGF-21 and Periostin were uniquely upregulated in female CFs treated with TGF-β1, whereas VEGF was uniquely upregulated in male CFs. TGF-β1 also uniquely downregulated LIX, Pentraxin 3, and MMP-3 in female CFs, and IGFBP6 and Lipocalin-2 in male CFs. Together, these data show that TGF-β1 activation of CFs on quiescence-supporting hydrogels elicits both shared and sex-specific secretory signatures. The overlap suggests a conserved myofibroblast activation cascade, whereas the sex-biased cytokines highlight distinct paracrine cues that may contribute to unique inflammatory responses in male versus female CFs in response to TGF-β1 (**Figure 6**).

**Fig. 6:**
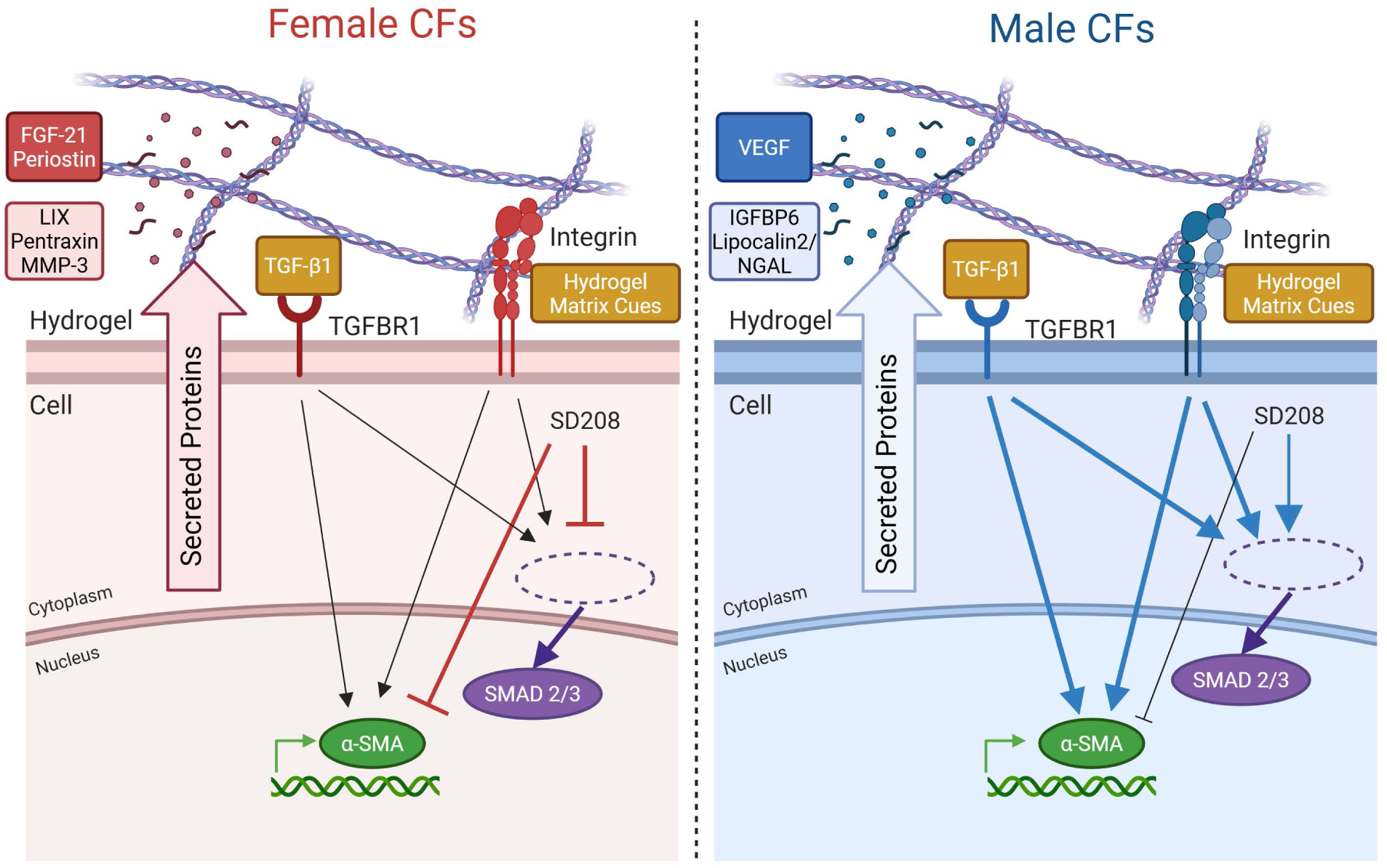
Proposed mechanisms for sex-specific cardiac myofibroblast activation cultured on hydrogels with TGF-β1. Thick lines indicate significant effect; thin lines indicate minor effect. Red lines indicate female biased effect, whereas blue lines indicate male biased effect. Arrow indicates activation, and bar indicates inhibition.

## Discussion

Here, we showed the effects of biological sex on TGF-β1 signaling in cardiac myofibroblasts and subsequent sex-dependent secretome alterations using engineered hydrogels. Our work describes for the first time to the best of our knowledge that hydrogels as cell culture platforms can successfully reveal sex differences in TGF-β1-treated cardiac myofibroblasts. Our contribution adds to a growing body of literature suggesting that hydrogels as cell culture platforms are essential tools to revealing sex-based differences in cellular phenotypes grown *in vitro*^54^. In line with previous work, hydrogels that recapitulate a healthy tissue matrix can retain fibroblast quiescence as opposed to TCPS, which has a supraphysiologic stiffness that automatically activates fibroblasts to myofibroblasts^55^. Our viscoelastic hydrogel formulation has a Young’s modulus of 24.1 ± 0.7 kPa, which mimics the mechanical properties of healthy myocardium measured with multiple techniques, including shear rheology (E ∼22 kPa) and uniaxial testing (E ∼5-50 kPa)^49,50^. Our synthetic PEG hydrogels represent one example of hydrogel design to recapitulate native myocardial tissue, which has also been used to support the culture of other cardiovascular cell types^56–60^. Recent work has used other synthetic and naturally derived materials to investigate sex differences in cellular phenotypes^40,41,45,46,61^. Beyond biomaterials-based tools, micro-physiological systems have also been employed to understand sex-based differences in human lung fibroblasts and endothelial cells exposed to steroidal sex hormones^62,63^. We anticipate the implementation of various bioengineered technologies will become increasingly useful to understand sex-based differences in cellular responses to the extracellular microenvironment.

Of note, we observed male CFs to be more responsive to TGF-β1 treatment relative to female CFs on hydrogels. TGF-β1 controls a myriad of intracellular biological processes including tissue development, cell proliferation, and cell migration in various cell types^64,65^. More specifically, previous work has identified that TGF-β1 directly regulates the expression of both α-SMA mRNA transcripts and α-SMA fibers to regulate cell structure and extracellular remodeling^66^. TGF-β1 has been observed to exert its effects through the regulation of SMAD2/3, which enters the nucleus after phosphorylation and directly induces α-SMA expression^67^. Previous studies treating male and female mouse CFs with TGF-β1 have reported no sex differences in myofibroblast activation on TCPS, although it was observed that male CFs and macrophages are more responsive to inflammatory factors like tumor necrosis factor alpha (TNF-α)^53^. Our study, which utilized hydrogels to maintain CF quiescence, demonstrates that male CFs are more responsive to TGF-β1 and exhibit increased myofibroblast activation in the presence of TGF-β1. We posit the sex difference in CF phenotype was exemplified using our engineered hydrogel system, highlighting the importance of using compliant biomaterials in the culture of mechanically sensitive cell types like fibroblasts.

Additionally, we observed female CFs to be more responsive to SD208 on hydrogels compared to male CFs. Previous studies have established SD208 as a small molecule drug that specifically targets TGF-β1 signaling via TGF-β1 receptor 1 kinase inhibition to prevent the activation of the transcription factors SMAD2/3 and subsequent myofibroblast activation^68^. Previous work has shown female porcine valvular interstitial cells require lower doses of SD208 than male cells to reduce activation by 20% compared to untreated samples^43^. In our work, we corroborate these findings by demonstrating that on hydrogels, female CFs showed significantly higher sensitivity to all concentrations of SD208 treatment compared to male CFs. Our work with establishing sex differences in cardiac myofibroblast responses to SD208 motivate future exploration into how other single drug and combination drug treatments (e.g., histone deacetylase inhibitors^69^) may impact cardiac myofibroblast phenotypes as a function of biological sex.

We also revealed sex differences in the CF secretome in response to TGF-β1, which may play a vital role in controlling autocrine and paracrine signaling in myocardial fibrosis. Crosstalk via secreted proteins and exosomes between cardiomyocytes, fibroblasts, inflammatory immune cells, and endothelial cells is known to orchestrate myocardial fibrosis progression^70,71^. TGF-β1 signaling is a critical pathway driving extracellular production of cytokines from CFs^72^. However, characterizations of how biological sex modulate secreted factors in response to inflammatory factors such as TGF-β1 have not been well described for CFs. One related study identified and isolated matrix-bound extracellular vesicles (EVs) from the left ventricle in humans and mice and determined microRNAs in EVs from aged males were pro-fibrotic, whereas microRNAs in EVs from both young and aged female myocardium showed anti-fibrotic effects in the heart^73^. Another study established the cardiac proteome is altered as a function of sex hormone and sex chromosome complement^74^. Here, we observed VEGF to be uniquely upregulated in male CFs, which may point to increased hypoxic stress and subsequent angiogenesis in fibrotic male hearts based on previous work^75,76^. We also observed FGF-21 to be uniquely upregulated in female CFs, which have previously been shown to be required for cardiac remodeling during pregnancy^77^. Periostin is also an essential driver of myocardial fibrosis and may play a role sex-dependent cardiac fibrosis in the aging mouse heart^78^. Together, we posit TGF-β1 signaling as a critical modulator of the secretome during sex-dependent myocardial fibrosis progression.

Our study has several limitations that warrant consideration in future work. First, although our work established baseline trends for male and female CF secretome within a physiologically relevant microenvironment, our study does not conclude whether TGF-β1-induced sex differences in gene expression and secretome are dependent on variations in sex hormone or sex chromosome biology. Future work can incorporate the Four Core Genotype (FCG) and XY* mouse models that have been genetically engineered to express phenotypic males and females with varied sex chromosome complement to decouple the relative contributions of sex chromosomes and gonadal sex hormones^79–81^. Other future work may include supplementing exogenous sex hormones into the CF culture media to detect relative sex differences associated with variations in sex hormone production that are associated with aging and may subsequently alter TGF-β1 signaling in myofibroblasts^62,82^. Second, our study relied on two-dimensional culture substrates to evaluate sex differences in cell phenotypes. We recognize that 3D and 4D culture strategies may enhance our spatial and temporal understanding of how male and female myofibroblasts cause sex differences in myocardial fibrosis in hydrogels of varied stiffness and viscoelasticity^37,38,83^. Furthermore, our study tested the effects of SD208, an experimental small molecule drug that has not been approved for use in humans. Future work may consider repurposing clinically approved inhibitors, including pirfenidone approved to treat symptoms of idiopathic pulmonary fibrosis^84^, to assess their efficacy in reducing cardiac myofibroblast activation.

In conclusion, our work demonstrates that TGF-β1 signaling, myofibroblast responses to SD208, and secretome composition are all impacted by biological sex, and that hydrogel culture enabled sex-dependent evaluations in cellular phenotypes. Our findings underscore the importance of integrating sex as a biological variable in preclinical myocardial fibrosis research and provide a steppingstone toward opportunities to identify sex-specific therapeutic interventions targeting sexually dimorphic signaling activity in cardiac myofibroblasts.

## Methods

### Hydrogel fabrication

Before hydrogel fabrication, 12 mm and 25 mm circular glass coverslips were treated with (3-mercaptopropyl)trimethoxysilane (MPTS, Cat. No. 175617, Sigma Aldrich) by vapor deposition at 60 °C overnight. The synthesis of poly(ethylene glycol) norbornene (PEG-Nb) was done following an established protocol. Hydrogel precursor solution composed of 20 kDa PEG-nb, 5 kDa PEG dithiol (Cat. No. A4075, Jenkem), cysteine-arginine-glycine-aspartic acid-serine (CRGDS, Bachem), and lithium phenyl-2,4,6-trimethylbenzoylphosphinate (LAP, Cat. No. 900889, Fisher Scientific) was mixed in the dark. The solution was then pipetted onto a Sigmacote (Cat. No. SL2, Sigma-Aldrich) treated glass slide and the circular coverslip was then lightly applied and pressed between the two surfaces to fully spread the solution over the coverslip. The slide was then cured under UV light at 10 mW/cm^2^ for 3 minutes. The slide was then immediately placed into a petri dish of PBS containing 5% isopropyl alcohol/PBS (IPA, Cat. No. A4591, Fisher Scientific) to sterilize and allow the hydrogel to release from the slide. A razor blade and forceps were used to remove the coverslips containing the hydrogel from the slide and placed them face up into a well plate. Hydrogels were then washed three times with PBS and allowed to swell overnight in CF media at 37°C and 5% CO2.

### Rheology

Rheology was conducted on photopolymerized hydrogels to measure the modulus of specific gel formulations. 30 μL of hydrogel precursor solution was pipetted onto a DHR3 (TA instruments) with a quartz base plate and 8 mm parallel plate geometry. The hydrogel solution was cured using an Omnicure set to 10 mW/cm^2^ for 3 min while the geometry exposed the gel solution to a constant 1% shear strain at 1 rad/s. The storage modulus (G’) and loss modulus (G”) were recorded for 1 minute before polymerization, 3 minutes during, and 1 minute after. The final storage modulus was then converted to Young’s modulus using the equation E = 2G′(1 + 𝛎), where 𝛎 is the Poisson’s ratio assumed to be 0.5 as G′>>>G″.

### Cardiac fibroblast isolation

CFs were obtained from 8-12-week-old adult C57BL/6 mice (Cat. No. #039108 Jackson Laboratories). For each isolation (i.e., biological replicate), ventricles from 3 hearts were pooled based on sex, and harvested cells were combined to generate one biological replicate for each sex. A total of 4 CF isolations per sex were used for this study to ensure biological reproducibility across experiments. Before each CF isolation, 6-well tissue culture plates (Cat. No. M8562, Cellvis) were coated with 1% gelatin (Cat. No. G9391, Sigma-Aldrich) for 1 hour and then sealed with Parafilm before being placed in a 4 °C fridge until use. On the day of isolation, gelatin-coated plates were warmed to 37 °C and thoroughly rinsed with phosphate-buffered saline (PBS).

After euthanasia, mouse hearts were removed and immediately placed into chilled Hank’s Balanced Salt Solution (HBSS, Cat. No. 14025076, Fisher Scientific) on ice. Hearts were vigorously squeezed with forceps to flush remaining blood out of the heart and then placed on a petri dish. The hearts were cut with forceps and a razor blade to remove the atria, and then the ventricles were quickly minced into homogenously small chunks. 5 mL of HBSS mixed with 4.375 μL DNase (Cat. No. 90083, Fisher Scientific), 125 μL Liberase TH (Cat. No. 540113500, Fisher Scientific), and 32.5 μL of 1M 4-(2-hydroxyethyl)-1-piperazineethanesulfonic acid (HEPES) were next added to assist with digestion. Hearts were minced again, mixed with 1 mL of digestion buffer, transferred to a conical tube, and placed on a plate shaker at 37 °C to further break down tissue components.

After 30 minutes of enzymatic digestion, the cell suspension was resuspended using a transfer pipette and allowed to shake for another 15 minutes. Afterwards, the suspension was immediately quenched with 25 mL of CF media [DMEM/F12 (Cat. No. 11320033, Thermo Scientific) basal media, 1 μg/mL amphotericin B (Cat. No. 15290026, Fisher Scientific), 50 U/mL penicillin and 50 μg/mL streptomycin (Cat. No. P4458, Sigma-Aldrich), and 30 μgi/mL of L-ascorbic acid (Cat. No. A5960, Sigma Aldrich)]. The digested tissue solution was then passed through a 100-μm cell strainer using a transfer pipette to remove pieces of decellularized tissue. The cell suspension was then centrifuged at 4 °C for 10 minutes at 400 g. The supernatant was then aspirated without disturbing the cell pellet and resuspended with 1X red blood cell lysis buffer (Cat. No. 00-4333-57, Thermo Scientific) for 5 minutes, then quenched with warmed CF media supplemented with 20% fetal bovine serum (FBS, Cat. No. 16000069, Thermo Scientific). The cell suspension was then centrifuged at 23 °C for 10 minutes at 400 g and the supernatant was aspirated as previously described. The cell pellet was then resuspended in 20% FBS CF media and distributed evenly into a gelatin-coated 6-well plate (cells from ∼1 heart were plated per well). The plate was then shaken gently to evenly distribute the cells in the well and then placed into the incubator at 37 °C for 4 hours to allow for the fibroblasts to selectively adhere to the plate.

### Cardiac fibroblast culture on hydrogels

After incubating for 4 hours, the wells were washed thoroughly with warmed PBS and cultured in 10% FBS CF media for up to 3 days to allow for expansion up to 80–90% confluency, changing media after 48-72 hours. After allowing the fibroblasts to expand, they were then collected using warm 1X trypsin (Cat. No. 15-400-054, Fisher Scientific) and centrifuged at 1000 rpm for 5 minutes. The cells were then counted using an automatic hemocytometer and seeded onto hydrogels or TCPS with designated conditioned media at 7,500 cells per well for immunostaining or 45,000 cells per well for real time quantitative polymerase chain reaction (RT-qPCR) and conditioned media retrieval experiments. CFs were freshly isolated and seeded at passage 1 (P1) for each experiment, as expansion for more than 4 days and multiple passages can induce CF senescence^37^. On the same day of seeding, male or female CFs were treated with recombinant human transforming growth factor beta 1 (TGF-β1, R&D Systems, Cat. No. 240-B) and/or SD208 (Selleck Chemicals, Cat. No. S7624) for 3 days prior to fixation.

### Immunostaining

Prior to fixing CFs for immunostaining, conditioned media was collected and stored at −20 °C before use. At specified time points, CFs were fixed with 4% paraformaldehyde (Cat. No. 50-980-487, Fisher Scientific) diluted in PBS for 20 minutes at room temperature (RT). After fixing, the paraformaldehyde was gently removed and cells were permeabilized with PBS containing 0.1% Triton-X 100 (Cat. No. AC215682500, Thermo Fisher Scientific) for 1 hour. Permeabilization buffer was removed and the cells were then blocked with PBS containing 5% bovine serum albumin (BSA, Cat. No. 50-550-391, Research Products International) for at least 1 hour or overnight at 4 °C. Immunostaining was performed with blocking buffer containing mouse anti-aSMA primary antibody (Cat. No. ab124964, Abcam) at 1:300 and rabbit anti-SMAD2/3 (Cat. No. ab202445, Abcam) primary antibody for 1 hour at RT. Primary antibody mixture was collected, and the CFs were then washed with PBS containing 0.05% Tween 20 (Cat. No. P1379, Sigma Aldrich) for 5 minutes on a shaker. After, samples were immersed in a secondary antibody solution of PBS containing donkey-anti-mouse Alexa Fluor 555 (Cat. No. A31570, Fisher Scientific) at 1:300, goat-anti-rabbit Alexa Fluor 488 (Cat. No. A11008, Fisher Scientific) at 1:300, 4′-6-diamidino-2-phenylindole (DAPI, Cat. No. 10236276001, Sigma Aldrich) at 1:500, and CellMask Deep Red stain (Cat. No. H32721, Fisher Scientific) at 1:5000 for 1 hour. Samples were then washed once with PBS containing 0.05% Tween 20 for 5 minutes on a shaker, followed by another wash with PBS. Finally, the samples were removed and placed glass side down in a glass-bottomed well plate and immersed in PBS at 4 °C until imaging. Imaging was performed using a Nikon Eclipse Ti2-E at 20X magnification and analyzed using an automated MATLAB script to quantify CF α-SMA gradient mean intensity, SMAD2/3 fluorescence intensity in the cytoplasm and the nucleus, and cell area.

### RNA isolation and real time quantitative polymerase chain reaction (RT-qPCR)

CFs were seeded at 45,000 cells per 25 mm hydrogel and cultured for 72 hours. RNA was isolated from the CFs using RNAeasy Micro Kit (Cat. No. 74004, Qiagen). Hydrogels were removed from culturing media and then inverted on top of the lysis buffer for at least 2 minutes. The hydrogel was then tilted and rinsed with an equal volume of 70% ethanol, and then collected into spin columns and purified according to RNAeasy Micro Kit manufacturer’s protocol to obtain total RNA. cDNA was synthesized using an iScript Synthesis kit (Cat. No. 1708841, Bio-Rad) to produce 100 ng of cDNA, which was stored at −20 °C until used. Relative gene expression of *FN*, *COL1A1*, and *ACTA2* were quantified using iQ SYBR Green Mix (Cat. No. 1708882, Bio-Rad) on a CFX 384 iCycler (Bio-Rad). Gene expression was normalized using the housekeeping gene B2M and compared against the male control values. All gene primer sequences are reported in **Supplemental Table 1**.

### Cytokine array

Conditioned media samples with secreted cytokines were collected three days after culture and pooled together from two technical replicates (one biological replicate). Conditioned media was then briefly centrifuged for 5 minutes at 1000 g to remove any suspended cellular debris from the proteins in the media. A Proteome Profiler Mouse XL Cytokine Array Kit (Cat. No. ARY028, Biotechne) was used to identify the expression of cytokines for each of the samples according to the manufacturer’s protocol. Briefly, membranes in the kit were used to capture the proteins using embedded antibodies arranged in a grid pattern, which are then easily identifiable using a template. Membranes are then closed in an autoradiography cassette with X-ray film to allow for the fluorescently tagged proteins to irradiate the film. Changes in protein expression were analyzed using the publicly available tool in Fiji called Protein Array Analyzer to create a grid structure in which protein abundance is quantified via fluorescence intensity. Analysis involved calculating the row-scaled value change of intensity values in inflammatory cytokine protein abundance in conditioned media samples from male or female CFs treated with TGF-β1 relative to sex-matched untreated controls.

### Statistical analysis

We analyzed immunostaining data using an Aligned Rank Transform (ART) nonparametric two-way with Tukey post-hoc tests or three-way ANOVA to test for main effects and interaction between factors, as implemented in the ARTool package in R and GraphPad PRISM. The threshold cutoff was set to P < 0.001. The Cohen’s d-value test was used in parallel to determine significant changes in the effect size to determine significance of means between two groups irrespective of large cell sample sizes. We analyzed cytokine array data by determining the fold change (Log2FC, cutoff ±1) and testing significance of relative cytokine abundance between two groups using Student t-tests (P<0.05). We analyzed RT-qPCR data using a two-way ANOVA (P < 0.05 with Tukey post-hoc tests) using GraphPad PRISM.

## Ethical Approval

Research was conducted in accordance with ethical guidelines at the University of California San Diego. *In vitro* use of mammalian cells was approved under Biological Use Authorization approval by the Institutional Biosafety Committee at the University of California San Diego. Animal studies were approved by the Institutional Animal Care and Use Committee (IACUC) Office at the University of California San Diego.

## Consent to Participate

No human studies were involved in this research.

## Consent to Publish

No human studies were involved in this research.

## Data Availability Statement

The data supporting the findings of this study are available upon reasonable request to the corresponding author.

## Acknowledgements

The authors would like to acknowledge Dr. Peng Guo and Dr. Richard Sanchez for their guidance on imaging at the UCSD Nikon Imaging Center.

## Author Contributions

M.F. and B.A.A conceived and supervised the study. M.F, A.K.N., R.M.G., and N.E.F.V. performed all experiments, data collection, and statistical analyses associated with this study. M.C.L. performed heart isolations and animal husbandry. M.F and B.A.A. created all figures and supplementary figures. M.F., A.K.N., and B.A.A wrote and edited the manuscript. All authors approved the manuscript.

## Funding

National Science Foundation Graduate Research Fellowship Program (NEFV)

National Institutes of Health, National Heart, Lung, and Blood Institute (NHLBI) F31 HL176153 (RMG)

National Institutes of Health Director’s New Innovator Award DP2 HL173948 (BAA)

Chan Zuckerberg Initiative Science Diversity Leadership Award DAF2022-309430 (BAA)

National Science Foundation CAREER 2442606 (BAA)

## References

1 Francis Stuart, S. D., De Jesus, N. M., Lindsey, M. L. & Ripplinger, C. M. The crossroads of inflammation, fibrosis, and arrhythmia following myocardial infarction. J Mol Cell Cardiol 91, 114–122 (2016). 10.1016/j.yjmcc.2015.12.024

2 Hinderer, S. & Schenke-Layland, K. Cardiac fibrosis – A short review of causes and therapeutic strategies. Adv Drug Deliver Rev 146, 77–82 (2019). 10.1016/j.addr.2019.05.011

3 Talman, V. & Ruskoaho, H. Cardiac fibrosis in myocardial infarction-from repair and remodeling to regeneration. Cell Tissue Res 365, 563–581 (2016). 10.1007/s00441-016-2431-9

4 Schimmel, K., Ichimura, K., Reddy, S., Haddad, F. & Spiekerkoetter, E. Cardiac Fibrosis in the Pressure Overloaded Left and Right Ventricle as a Therapeutic Target. Front Cardiovasc Med 9, 886553 (2022). 10.3389/fcvm.2022.886553

5 Travers, J. G., Tharp, C. A., Rubino, M. & McKinsey, T. A. Therapeutic targets for cardiac fibrosis: from old school to next-gen. The Journal of Clinical Investigation 132 (2022). 10.1172/JCI148554

6 Zhang, H. et al. Multiscale drug screening for cardiac fibrosis identifies MD2 as a therapeutic target. Cell 187, 7143–7163.e7122 (2024). 10.1016/j.cell.2024.09.034

7 Martin, T. G. & Leinwand, L. A. Hearts apart: sex differences in cardiac remodeling in health and disease. J Clin Invest 134 (2024). 10.1172/jci180074

8 Regitz-Zagrosek, V. & Kararigas, G. Mechanistic Pathways of Sex Differences in Cardiovascular Disease. Physiological Reviews 97, 1–37 (2016). 10.1152/physrev.00021.2015

9 Collins, H. E. Female cardiovascular biology and resilience in the setting of physiological and pathological stress. Redox Biology 63, 102747 (2023). 10.1016/j.redox.2023.102747

10 Leinwand, L. A. & Crocini, C. Age-Dependent Mechanisms of Cardiac Hypertrophy Regression Following Exercise in Female Mice. bioRxiv, 2025.2004.2007.647563 (2025). 10.1101/2025.04.07.647563

11 Scantlebury, D. C. & Borlaug, B. A. Why are women more likely than men to develop heart failure with preserved ejection fraction? Curr Opin Cardiol 26, 562–568 (2011). 10.1097/HCO.0b013e32834b7faf

12 Duca, F. et al. Gender-related differences in heart failure with preserved ejection fraction. Scientific Reports 8, 1080 (2018). 10.1038/s41598-018-19507-7

13 Cavasin, M. A., Tao, Z., Menon, S. & Yang, X. P. Gender differences in cardiac function during early remodeling after acute myocardial infarction in mice. Life Sci 75, 2181–2192 (2004). 10.1016/j.lfs.2004.04.024

14 Haines, C. D., Harvey, P. A. & Leinwand, L. A. Estrogens mediate cardiac hypertrophy in a stimulus-dependent manner. Endocrinology 153, 4480–4490 (2012). 10.1210/en.2012-1353

15 Bauer, M., Gadkari, M., Martinez Yus, M., Santhanam, L. & Steppan, J. Sexual dimorphism in animal models of heart failure with preserved ejection fraction. J Appl Physiol (1985) 138, 1449–1473 (2025). 10.1152/japplphysiol.00595.2024

16 Walker, C. J., Schroeder, M. E., Aguado, B. A., Anseth, K. S. & Leinwand, L. A. Matters of the heart: Cellular sex differences. J Mol Cell Cardiol 160, 42–55 (2021). 10.1016/j.yjmcc.2021.04.010

17 Tallquist, M. D. & Molkentin, J. D. Redefining the identity of cardiac fibroblasts. Nature Reviews Cardiology 14, 484–491 (2017). 10.1038/nrcardio.2017.57

18 Souders, C. A., Bowers, S. L. K. & Baudino, T. A. Cardiac Fibroblast. Circulation Research 105, 1164–1176 (2009). 10.1161/CIRCRESAHA.109.209809

19 Travers, J. G., Kamal, F. A., Robbins, J., Yutzey, K. E. & Blaxall, B. C. Cardiac Fibrosis: The Fibroblast Awakens. Circ Res 118, 1021–1040 (2016). 10.1161/circresaha.115.306565

20 Peter, A. K. et al. Cardiac Fibroblasts Mediate a Sexually Dimorphic Fibrotic Response to β-Adrenergic Stimulation. J Am Heart Assoc 10, e018876 (2021). 10.1161/jaha.120.018876

21 Medzikovic, L., Aryan, L. & Eghbali, M. Connecting sex differences, estrogen signaling, and microRNAs in cardiac fibrosis. J Mol Med (Berl) 97, 1385–1398 (2019). 10.1007/s00109-019-01833-6

22 Ramli, M. F. H., Aguado, B. A. & Young, J. L. Signals from the extracellular matrix: Region- and sex-specificity in cardiac aging. Current Opinion in Cell Biology 95, 102524 (2025). 10.1016/j.ceb.2025.102524

23 Rurik, J. G., Aghajanian, H. & Epstein, J. A. Immune Cells and Immunotherapy for Cardiac Injury and Repair. Circ Res 128, 1766–1779 (2021). 10.1161/circresaha.121.318005

24 Bengel, F. M., Hess, A., Diekmann, J. & Thackeray, J. T. Molecular imaging of immune and fibrosis targets to guide therapy for repair after myocardial infarction. Nature Reviews Cardiology (2026). 10.1038/s41569-025-01242-y

25 Shimazaki, M. et al. Periostin is essential for cardiac healing after acute myocardial infarction. J Exp Med 205, 295–303 (2008). 10.1084/jem.20071297

26 Aguado, B. A. et al. Transcatheter aortic valve replacements alter circulating serum factors to mediate myofibroblast deactivation. Sci Transl Med 11 (2019). 10.1126/scitranslmed.aav3233

27 Frangogiannis, N. Transforming growth factor-β in tissue fibrosis. J Exp Med 217, e20190103 (2020). 10.1084/jem.20190103

28 Shinde, A. V., Humeres, C. & Frangogiannis, N. G. The role of α-smooth muscle actin in fibroblast-mediated matrix contraction and remodeling. Biochimica et Biophysica Acta (BBA) - Molecular Basis of Disease 1863, 298–309 (2017). 10.1016/j.bbadis.2016.11.006

29 Weber, K. T., Sun, Y., Bhattacharya, S. K., Ahokas, R. A. & Gerling, I. C. Myofibroblast-mediated mechanisms of pathological remodelling of the heart. Nat Rev Cardiol 10, 15–26 (2013). 10.1038/nrcardio.2012.158

30 Bahr, J. et al. A secretome atlas of cardiac fibroblasts from healthy and infarcted mouse hearts. Communications Biology 8, 675 (2025). 10.1038/s42003-025-08083-y

31 Bräuninger, H. et al. Cytokine-Mediated Alterations of Human Cardiac Fibroblast’s Secretome. International Journal of Molecular Sciences 22, 12262 (2021).

32 Ceccato, T. L. et al. Defining the Cardiac Fibroblast Secretome in a Fibrotic Microenvironment. Journal of the American Heart Association 9, e017025 (2020). 10.1161/JAHA.120.017025

33 Montalvo, C. et al. Androgens Contribute to Sex Differences in Myocardial Remodeling under Pressure Overload by a Mechanism Involving TGF-β. PLOS ONE 7, e35635 (2012). 10.1371/journal.pone.0035635

34 Vogt, B. J. et al. Determining sex differences in aortic valve myofibroblast responses to drug combinations identified using a digital medicine platform. Science Advances 11, eadu2695 10.1126/sciadv.adu2695

35 Nagaraju Chandan, K., et al. Myofibroblast Phenotype and Reversibility of Fibrosis in Patients With End-Stage Heart Failure. JACC 73, 2267–2282 (2019). 10.1016/j.jacc.2019.02.049

36 Cho, S. et al. Selective inhibition of stromal mechanosensing suppresses cardiac fibrosis. Nature 642, 766–775 (2025). 10.1038/s41586-025-08945-9

37 Felisbino, M. B. et al. Substrate stiffness modulates cardiac fibroblast activation, senescence, and proinflammatory secretory phenotype. Am J Physiol Heart Circ Physiol 326, H61–h73 (2024). 10.1152/ajpheart.00483.2023

38 Herum, K. M., Choppe, J., Kumar, A., Engler, A. J. & McCulloch, A. D. Mechanical regulation of cardiac fibroblast profibrotic phenotypes. Mol Biol Cell 28, 1871–1882 (2017). 10.1091/mbc.E17-01-0014

39 Caliari, S. R. & Burdick, J. A. A practical guide to hydrogels for cell culture. Nature Methods 13, 405–414 (2016). 10.1038/nmeth.3839

40 Aguado, B. A. et al. Genes That Escape X Chromosome Inactivation Modulate Sex Differences in Valve Myofibroblasts. Circulation 145, 513–530 (2022). 10.1161/CIRCULATIONAHA.121.054108

41 Gorashi, R. M. et al. Y chromosome–linked UTY modulates sex differences in valvular fibroblast methylation in response to nanoscale extracellular matrix cues. Science Advances 11, eads5717 (2025). doi:10.1126/sciadv.ads5717

42 Vogt, B. J., Peters, D. K., Anseth, K. S. & Aguado, B. A. Inflammatory serum factors from aortic valve stenosis patients modulate sex differences in valvular myofibroblast activation and osteoblast-like differentiation. Biomater. Sci. 10, 6341–6353 (2022). 10.1039/D2BM00844K

43 Vogt, B. J. et al. Determining sex differences in aortic valve myofibroblast responses to drug combinations identified using a digital medicine platform. Science Advances 11, eadu2695 (2025). doi:10.1126/sciadv.adu2695

44 Félix Vélez, N. E., Tu, K., Guo, P., Reeves, R. R. & Aguado, B. A. Secreted Cytokines From Inflammatory Macrophages Modulate Sex Differences in Valvular Interstitial Cells on Hydrogel Biomaterials. Journal of Biomedical Materials Research Part A 113, e37885 (2025). 10.1002/jbm.a.37885

45 Nkansah, A. et al. Characterization of Sex-Based Differences in Integrin-Mediated Endothelial Cell Adhesion to Bioactive Hydrogels. ACS Biomaterials Science & Engineering 11, 6515–6520 (2025). 10.1021/acsbiomaterials.5c01461

46 James, B. D. & Allen, J. B. Sex-Specific Response to Combinations of Shear Stress and Substrate Stiffness by Endothelial Cells In Vitro. Advanced Healthcare Materials 10, 2100735 (2021). 10.1002/adhm.202100735

47 Félix Vélez, N. E., Gorashi, R. M. & Aguado, B. A. Chemical and molecular tools to probe biological sex differences at multiple length scales. Journal of Materials Chemistry B 10, 7089–7098 (2022). 10.1039/D2TB00871H

48 Aguado, B. A., Grim, J. C., Rosales, A. M., Watson-Capps, J. J. & Anseth, K. S. Engineering precision biomaterials for personalized medicine. Sci Transl Med 10, eaam8645 (2018). 10.1126/scitranslmed.aam8645

49 Emig, R. et al. Passive myocardial mechanical properties: meaning, measurement, models. Biophys Rev 13, 587–610 (2021). 10.1007/s12551-021-00838-1

50 Guimarães, C. F., Gasperini, L., Marques, A. P. & Reis, R. L. The stiffness of living tissues and its implications for tissue engineering. Nature Reviews Materials 5, 351–370 (2020). 10.1038/s41578-019-0169-1

51 Fairbanks, B. D. et al. A Versatile Synthetic Extracellular Matrix Mimic via Thiol-Norbornene Photopolymerization. Adv Mater 21, 5005–5010 (2009). 10.1002/adma.200901808

52 Mancini, D. et al. New methodologies to accurately assess circulating active transforming growth factor-β1 levels: implications for evaluating heart failure and the impact of left ventricular assist devices. Translational Research 192, 15–29 (2018). 10.1016/j.trsl.2017.10.006

53 Barcena, M. L. et al. Male Macrophages and Fibroblasts from C57/BL6J Mice Are More Susceptible to Inflammatory Stimuli. Front Immunol 12, 758767 (2021). 10.3389/fimmu.2021.758767

54 Fogg, K. et al. Roadmap on biomaterials for women’s health. Journal of Physics: Materials 6, 012501 (2023). 10.1088/2515-7639/ac90ee

55 Wang, H., Tibbitt, M. W., Langer, S. J., Leinwand, L. A. & Anseth, K. S. Hydrogels preserve native phenotypes of valvular fibroblasts through an elasticity-regulated PI3K/AKT pathway. Proc Natl Acad Sci U S A 110, 19336–19341 (2013). 10.1073/pnas.1306369110

56 Puperi, D. S., Balaoing, L. R., O’Connell, R. W., West, J. L. & Grande-Allen, K. J. 3-Dimensional spatially organized PEG-based hydrogels for an aortic valve co-culture model. Biomaterials 67, 354–364 (2015). 10.1016/j.biomaterials.2015.07.039

57 Kang, L. H. et al. Optimizing Photo-Encapsulation Viability of Heart Valve Cell Types in 3D Printable Composite Hydrogels. Ann Biomed Eng 45, 360–377 (2017). 10.1007/s10439-016-1619-1

58 Amitrano, M. J., Cho, M., Coughlin, E. M., Palecek, S. P. & Murphy, W. L. Synthetic hydrogels support robust and reproducible cardiomyocyte differentiation. Biomater. Sci. 13, 2142–2151 (2025). 10.1039/D4BM01636J

59 Wu, Y., Jane Grande-Allen, K. & West, J. L. Adhesive Peptide Sequences Regulate Valve Interstitial Cell Adhesion, Phenotype and Extracellular Matrix Deposition. Cellular and Molecular Bioengineering 9, 479–495 (2016). 10.1007/s12195-016-0451-x

60 Bretherton, R. C. et al. Preventing hypocontractility-induced fibroblast expansion alleviates dilated cardiomyopathy. Science 390, eadv9157 10.1126/science.adv9157

61 Simon, L. R., Scott, A. J., Figueroa Rios, L., Zembles, J. & Masters, K. S. Cellular-scale sex differences in extracellular matrix remodeling by valvular interstitial cells. Heart Vessels 38, 122–130 (2023). 10.1007/s00380-022-02164-2

62 Martier, A. T. et al. Estradiol and dihydrotestosterone exert sex-specific effects on human fibroblast and endothelial proliferation, bioenergetics, and vasculogenesis. Communications Biology 8, 1422 (2025). 10.1038/s42003-025-08822-1

63 Martier, A., Wills Kpeli, G., Boone, K., Posey, I. R. & Mondrinos, M. J. Fetal Bovine Serum Modulates Primary Human Cell Phenotypes, Endothelial Barrier Function, Vasculogenesis, and Angiogenesis in a Sex-Specific Manner. Cellular and Molecular Bioengineering 18, 433–449 (2025). 10.1007/s12195-025-00860-3

64 Massagué, J. & Sheppard, D. TGF-β signaling in health and disease. Cell 186, 4007–4037 (2023). 10.1016/j.cell.2023.07.036

65 Hinz, B. The extracellular matrix and transforming growth factor-β1: Tale of a strained relationship. Matrix Biology 47, 54–65 (2015). 10.1016/j.matbio.2015.05.006

66 Roy, S. G., Nozaki, Y. & Phan, S. H. Regulation of alpha-smooth muscle actin gene expression in myofibroblast differentiation from rat lung fibroblasts. Int J Biochem Cell Biol 33, 723–734 (2001). 10.1016/s1357-2725(01)00041-3

67 Khalil, H. et al. Fibroblast-specific TGF-β-Smad2/3 signaling underlies cardiac fibrosis. J Clin Invest 127, 3770–3783 (2017). 10.1172/jci94753

68 Uhl, M. et al. SD-208, a novel transforming growth factor beta receptor I kinase inhibitor, inhibits growth and invasiveness and enhances immunogenicity of murine and human glioma cells in vitro and in vivo. Cancer Res 64, 7954–7961 (2004). 10.1158/0008-5472.Can-04-1013

69 McKinsey, T. A. Therapeutic potential for HDAC inhibitors in the heart. Annu Rev Pharmacol Toxicol 52, 303–319 (2012). 10.1146/annurev-pharmtox-010611-134712

70 Li, Y., Asfour, H. & Bursac, N. Age-dependent functional crosstalk between cardiac fibroblasts and cardiomyocytes in a 3D engineered cardiac tissue. Acta Biomaterialia 55, 120–130 (2017). 10.1016/j.actbio.2017.04.027

71 Bowers, S. L. K., Meng, Q. & Molkentin, J. D. Fibroblasts orchestrate cellular crosstalk in the heart through the ECM. Nature Cardiovascular Research 1, 312–321 (2022). 10.1038/s44161-022-00043-7

72 Chowkwale, M., Lindsey, M. L. & Saucerman, J. J. Intercellular model predicts mechanisms of inflammation-fibrosis coupling after myocardial infarction. J Physiol 601, 2635–2654 (2023). 10.1113/jp283346

73 Ronan, G., Bahcecioglu, G., Yang, J. & Zorlutuna, P. Cardiac tissue-resident vesicles differentially modulate anti-fibrotic phenotype by age and sex through synergistic miRNA effects. Biomaterials 311, 122671 (2024). 10.1016/j.biomaterials.2024.122671

74 Shi, W. et al. Cardiac proteomics reveals sex chromosome-dependent differences between males and females that arise prior to gonad formation. Dev Cell 56, 3019–3034.e3017 (2021). 10.1016/j.devcel.2021.09.022

75 Räsänen, M. et al. VEGF-B Promotes Endocardium-Derived Coronary Vessel Development and Cardiac Regeneration. Circulation 143, 65–77 (2021). 10.1161/CIRCULATIONAHA.120.050635

76 Shen, J. et al. Increased myocardial stiffness activates cardiac microvascular endothelial cell via VEGF paracrine signaling in cardiac hypertrophy. J Mol Cell Cardiol 122, 140–151 (2018). 10.1016/j.yjmcc.2018.08.014

77 Redondo-Angulo, I. et al. Fgf21 is required for cardiac remodeling in pregnancy. Cardiovasc Res 113, 1574–1584 (2017). 10.1093/cvr/cvx088

78 Angelini, A. et al. Sex-specific phenotypes in the aging mouse heart and consequences for chronic fibrosis. American Journal of Physiology-Heart and Circulatory Physiology 323, H285–H300 (2022). 10.1152/ajpheart.00078.2022

79 Shi, W. et al. Cardiac proteomics reveals sex chromosome-dependent differences between males and females that arise prior to gonad formation. Developmental Cell 56, 3019–3034.e3017 (2021). 10.1016/j.devcel.2021.09.022

80 Reue, K. & Wiese, C. B. Illuminating the Mechanisms Underlying Sex Differences in Cardiovascular Disease. Circulation Research 130, 1747–1762 (2022). 10.1161/CIRCRESAHA.122.320259

81 Arnold, A. P. & Chen, X. What does the “four core genotypes” mouse model tell us about sex differences in the brain and other tissues? Front Neuroendocrinol 30, 1–9 (2009). 10.1016/j.yfrne.2008.11.001

82 Watts, K. & Richardson, W. J. Effects of Sex and 17 β-Estradiol on Cardiac Fibroblast Morphology and Signaling Activities In Vitro. Cells 10, 2564 (2021).

83 Saini, H., Navaei, A., Van Putten, A. & Nikkhah, M. 3D Cardiac Microtissues Encapsulated with the Co-Culture of Cardiomyocytes and Cardiac Fibroblasts. Advanced Healthcare Materials 4, 1961–1971 (2015). 10.1002/adhm.201500331

84 Lewis, G. A. et al. Pirfenidone in heart failure with preserved ejection fraction: a randomized phase 2 trial. Nature Medicine 27, 1477–1482 (2021). 10.1038/s41591-021-01452-0

