## Supplementary Materials for "Transforming Growth Factor β1 Modulates Sex Differences in Cardiac Myofibroblast Activation on Hydrogel Biomaterials"

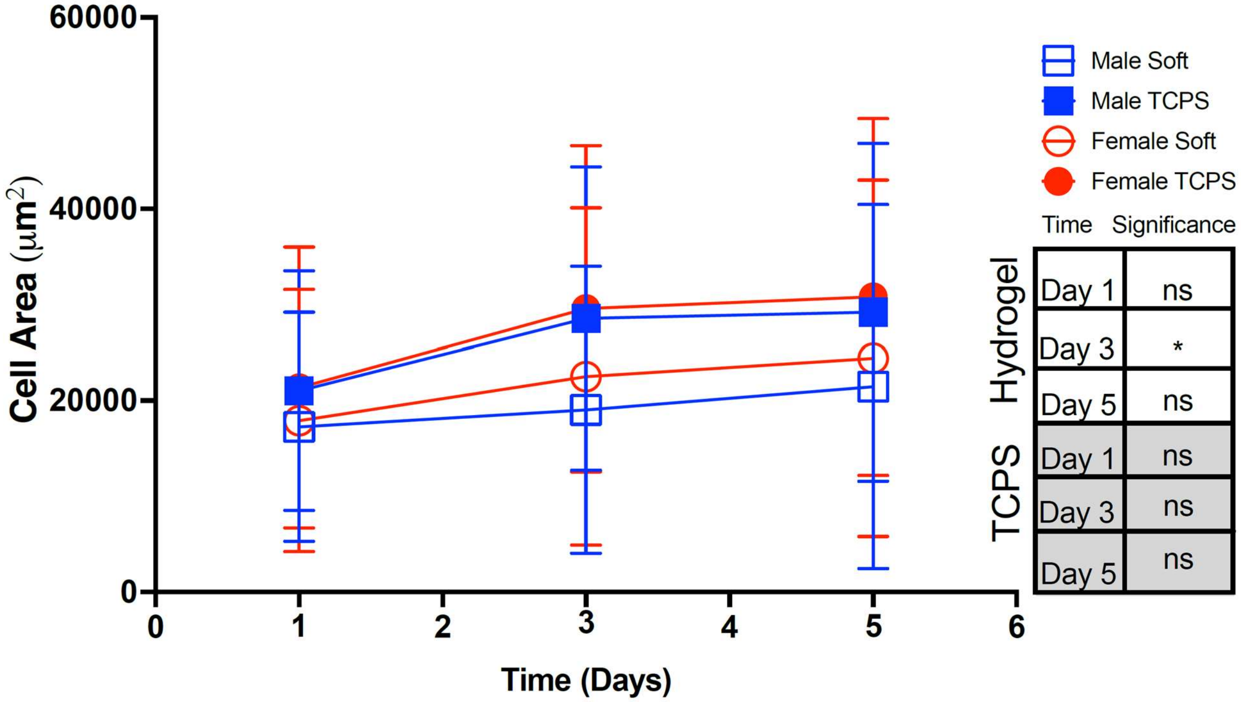


**Supplementary Figure 1: Cell area of cardiac fibroblasts cultured on hydrogels or TCPS.** Quantification of cell area in male and female CFs cultured on hydrogels and TCPS at various time points. N=3 hydrogels, mean ± S.D. shown. Significance was determined using two-way nonparametric ANOVA ($P < 0.05$) and a Cohen's d test (*$d < 0.2$), denoting significance between sex in the table provided.

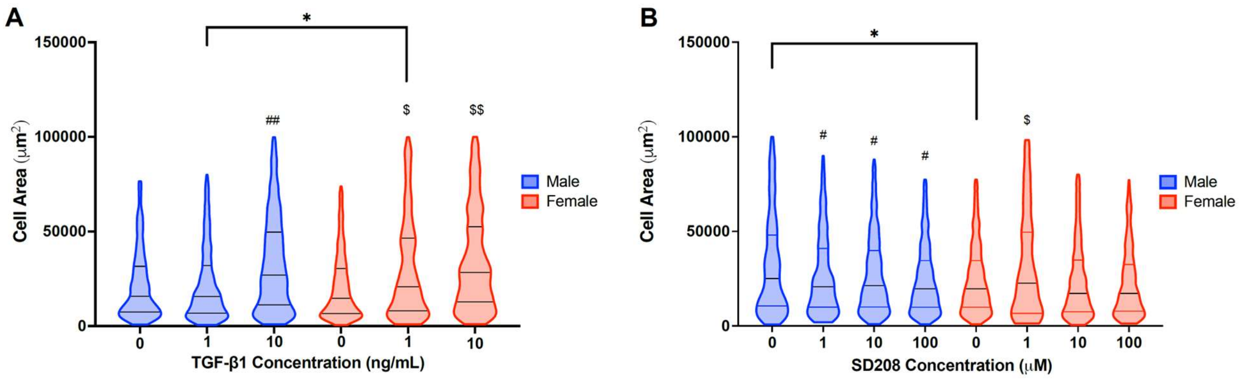


**Supplementary Figure 2: Cell area for cardiac fibroblasts treated with TGFβ-1or SD208.** Quantification of cell area in male and female CFs cultured on hydrogels and TCPS when treated with either (A) 0, 1, and 10 ng/mL TGFβ-1 or (B) with 0, 1, 10, and 100 μM of SD208 . N=3 hydrogels, n>432 cells, mean ± S.D. shown. Significance was determined using two-way nonparametric ANOVA (P  < 0.05) and a Cohen's d test (*d < 0.2, **d < 0.5, ***d < 0.8, and ****d < 1.4) denoting significance between sex; asterisk indicates significance in male group ($^{\#}$d < 0.2, $^{\#\#}$d < 0.5, $^{\#\#\#}$d < 0.8, and $^{\#\#\#\#}$d < 1.4) denoting significance relative to male control;  dollar sign indicates significance in female group ($^{\$}$d < 0.2, $^{\$\$}$d < 0.5, $^{\$\$\$}$d < 0.8, and $^{\$\$\$\$}$d < 1.4) denoting significance relative to female control.

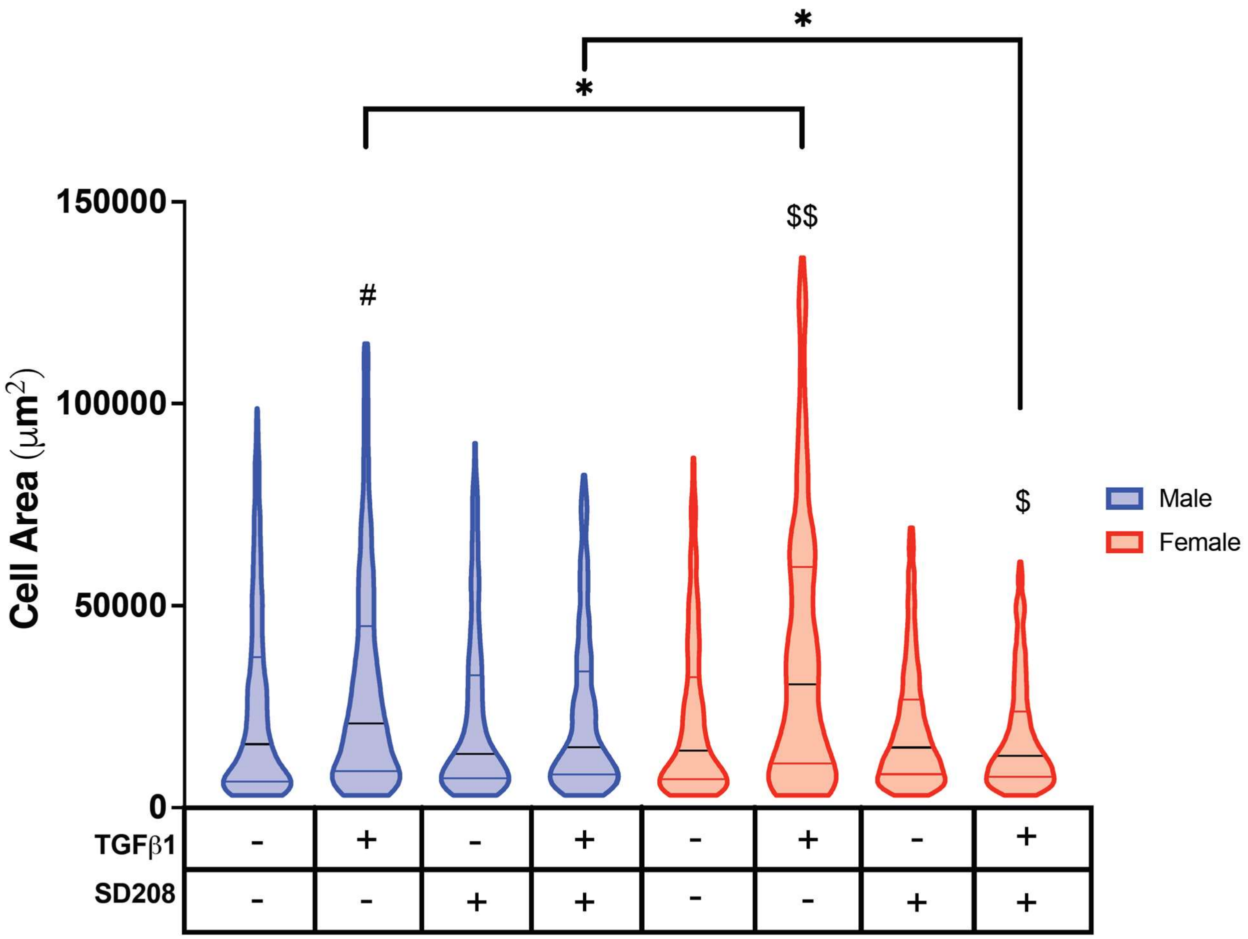


**Supplementary Figure 3: Cell area for cardiac fibroblasts treated with combined media.** Quantification of cell area for male and female cardiac fibroblasts cultured on hydrogels TGF-β1 and SD208. N=3 hydrogels, mean ± S.D. shown. Significance was determined using two-way nonparametric ANOVA ($P < 0.05$) and a Cohen's d test (*d < 0.2, **d < 0.5, ***d < 0.8, and ****d < 1.4) denoting significance between sex; asterisk indicates significance in male group ($^{\#}$d < 0.2, $^{\#\#}$d < 0.5, $^{\#\#\#}$d < 0.8, and $^{\#\#\#\#}$d < 1.4) denoting significance relative to male control; dollar sign indicates significance in female group ($^{\$}$d < 0.2, $^{\$\$}$d < 0.5, $^{\$\$\$}$d < 0.8, and $^{\$\$\$\$}$d < 1.4) denoting significance relative to female control.

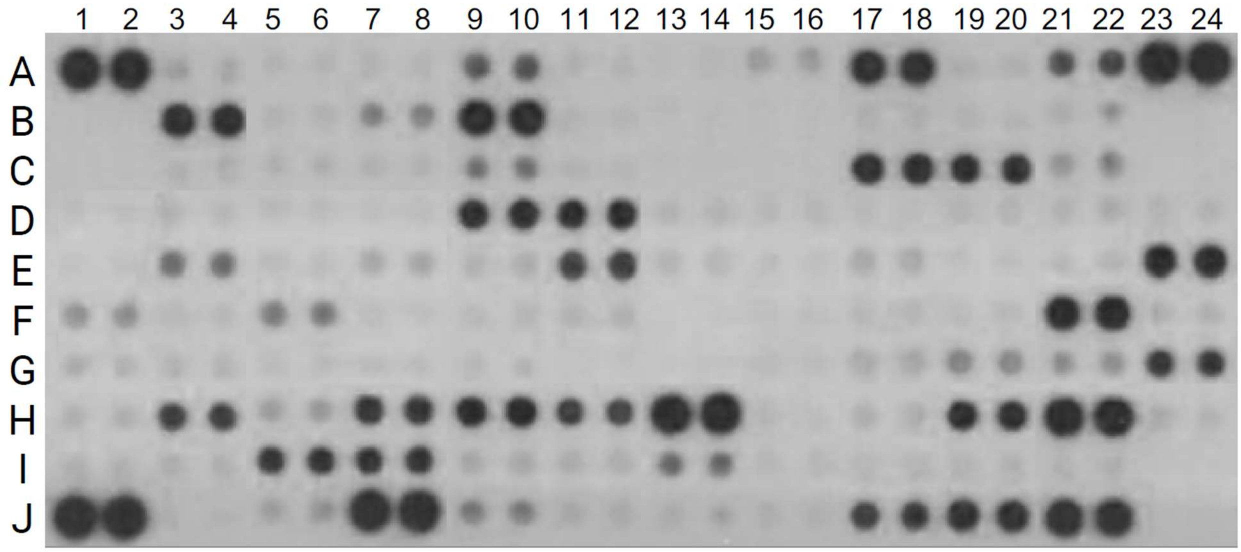


**Supplementary Figure 4: Sample Cytokine Array Blot.** Row/column IDs for each cytokine spot is provided in Supplementary Table 2.

**Supplementary Table 1: RT-qPCR Primer Sequences**

| Gene Name | Primer Sequence | Direction |
|---|---|---|
| ***B2M*** | GGTCTTTCTGGTGCTTGTCT | Forward |
| | ACGTAGCAGTTCAGTATGTTCG | Reverse |
| ***COL1A1*** | TGGTGAAGCAGGCAAGC | Forward |
| | GAAACCTCTCTCGCCTCTTG | Reverse |
| ***FN*** | GAGCTATCCATTTCACCTTCAGA | Forward |
| | TTGTTCGTAGACACTGGAGAC | Reverse |
| ***ACTA2*** | TCAGCGTTCAGCCTCC | Forward |
| | CCAGAGCCATTGTCGC | Reverse |

**Supplementary Table 2: Cytokine Array Protein Targets**

| Spot(s) | Protein Name | Spot(s) | Protein Name | Spot(s) | Protein Name |
|---|---|---|---|---|---|
| A1, A2 | Reference Spots | D13, D14 | DKK-1 | G17, G18 | IL-27 p28 |
| A3, A4 | Adiponectin/Acrp30 | D15, D16 | DPPIV/CD26 | G19, G20 | IL-28A/B |
| A5, A6 | Amphiregulin | D17, D18 | EGF | G21, G22 | IL-33 |
| A7, A8 | Angiopoietin-1 | D19, D20 | Endoglin/CD105 | G23, G24 | LDL R |
| A9, A10 | Angiopoietin-2 | D21, D22 | Endostatin | H1, H2 | Leptin |
| A11, A12 | Angiopoietin-like 3 | D23, D24 | Fetuin A/AHSG | H3, H4 | LIF |
| A13, A14 | BAFF/BLyS/TNFSF13B | E1, E2 | FGF acidic | H5, H6 | Lipocalin-2/NGAL |
| A15, A16 | C1q R1/CD93 | E3, E4 | FGF-21 | H7, H8 | LIX |
| A17, A18 | CCL2/JE/MCP-1 | E5, E6 | Flt-3 Ligand | H9, H10 | M-CSF |
| A19, A20 | CCL3/CCL4/MIP-1α/β | E7, E8 | Gas 6 | H11, H12 | MMP-2 |
| A21, A22 | CCL5/RANTES | E9, E10 | G-CSF | H13, H14 | MMP-3 |
| A23, A24 | Reference Spots | E11, E12 | GDF-15 | H15, H16 | MMP-9 |
| B3, B4 | CCL6/C10 | E13, E14 | GM-CSF | H17, H18 | Myeloperoxidase |
| B5, B6 | CCL11/Eotaxin | E15, E16 | HGF | H19, H20 | Osteopontin (OPN) |
| B7, B8 | CCL12/MCP-5 | E17, E18 | ICAM-1/CD54 | H21, H22 | Osteoprotegerin/TNFRSF11B |
| B9, B10 | CCL17/TARC | E19, E20 | IFN-γ | H23, H24 | PD-ECGF/Thymidine phosphorylase |
| B11, B12 | CCL19/MIP-3β | E21, E22 | IGFBP-1 | I1, I2 | PDGF-BB |
| B13, B14 | CCL20/MIP-3α | E23, E24 | IGFBP-2 | I3, I4 | Pentraxin 2/SAP |
| B15, B16 | CCL21/6Ckine | F1, F2 | IGFBP-3 | I5, I6 | Pentraxin 3/TSG-14 |
| B17, B18 | CCL22/MDC | F3, F4 | IGFBP-5 | I7, I8 | Periostin/OSF-2 |
| B19, B20 | CD14 | F5, F6 | IGFBP-6 | I9, I10 | Pref-1/DLK-1/FA1 |
| B21, B22 | CD40/TNFRSF5 | F7, F8 | IL-1α/IL-1F1 | I11, I12 | Proliferin |
| C3, C4 | CD160 | F9, F10 | IL-1β/IL-1F2 | I13, I14 | Proprotein Convertase 9/PCSK9 |
| C5, C6 | Chemerin | F11, F12 | IL-1ra/IL-1F3 | I15, I16 | RAGE |
| C7, C8 | Chitinase 3-like 1 | F13, F14 | IL-2 | I17, I18 | RBP4 |

| | | | | | |
|---|---|---|---|---|---|
| C9, C10 | Coagulation Factor III/Tissue Factor | F15, F16 | IL-3 | I19, I20 | Reg3G |
| C11, C12 | Complement Component C5/C5a | F17, F18 | IL-4 | I21, I22 | Resistin |
| C13, C14 | Complement Factor D | F19, F20 | IL-5 | J1, J2 | Reference Spots |
| C15, C16 | C-Reactive Protein/CRP | F21, F22 | IL-6 | J3, J4 | E-Selectin/CD62E |
| C17, C18 | CX3CL1/Fractalkine | F23, F24 | IL-7 | J5, J6 | P-Selectin/CD62P |
| C19, C20 | CXCL1/KC | G1, G2 | IL-10 | J7, J8 | Serpin E1/PAI-1 |
| C21, C22 | CXCL2/MIP-2 | G3, G4 | IL-11 | J9, J10 | Serpin F1/PEDF |
| D1, D2 | CXCL9/MIG | G5, G6 | IL-12 p40 | J11, J12 | Thrombopoietin |
| D3, D4 | CXCL10/IP-10 | G7, G8 | IL-13 | J13, J14 | TIM-1/KIM-1/HAVCR |
| D5, D6 | CXCL11/I-TAC | G9, G10 | IL-15 | J15, J16 | TNF-α |
| D7, D8 | CXCL13/BLC/BCA-1 | G11, G12 | IL-17A | J17, J18 | VCAM-1/CD106 |
| D9, D10 | CXCL16 | G13, G14 | IL-22 | J19, J20 | VEGF |
| D11, D12 | Cystatin C | G15, G16 | IL-23 | J21, J22 | WISP-1/CCN4 |
